# RNA-Binding Proteins Direct Myogenic Cell Fate Decisions

**DOI:** 10.1101/2021.03.14.435333

**Authors:** Joshua R. Wheeler, Oscar N. Whitney, Thomas O. Vogler, Eric D. Nguyen, Bradley Pawlikowski, Evan Lester, Alicia Cutler, Tiffany Elston, Nicole Dalla Betta, Kevin R. Parker, Kathryn E. Yost, Hannes Vogel, Thomas A. Rando, Howard Y. Chang, Aaron M. Johnson, Roy Parker, Bradley B. Olwin

## Abstract

RNA-binding proteins (RBPs) are essential for skeletal muscle regeneration and RBP dysfunction causes muscle degeneration and neuromuscular disease. How ubiquitously expressed RBPs orchestrate complex tissue regeneration and direct cell fate decisions in skeletal muscle remains poorly understood. Single cell RNA-sequencing of regenerating skeletal muscle reveals that RBP expression, including numerous neuromuscular disease-associated RBPs, is temporally regulated in skeletal muscle stem cells and correlates to stages of myogenic differentiation. By combining machine learning with RBP engagement scoring, we discover that the neuromuscular disease associated RBP Hnrnpa2b1 is a differentiation-specifying regulator of myogenesis controlling myogenic cell fate transitions during terminal differentiation. The timing of RBP expression specifies cell fate transitions by providing a layer of post-transcriptional regulation needed to coordinate stem cell fate decisions during complex tissue regeneration.

## INTRODUCTION

Skeletal muscle is among the longest-lived tissues in the human body, is essential for locomotion, respiration, and longevity, and thus requires constant maintenance (Sharples et al., 2015). Skeletal muscle is comprised of post-mitotic myofibers that house resident muscle stem cells (MuSCs), which have the remarkable ability to repair skeletal muscle following damage (Lepper et al., 2011; Murphy et al., 2011; Sambasivan et al., 2011; Shi and Garry, 2006). MuSCs typically reside in a quiescent state (Feige et al., 2018). However, in response to muscle injury, MuSCs become activated, expand and then either self-renew or differentiate into progenitors capable of repairing myofibers (Baghdadi and Tajbakhsh, 2018; Dumont et al., 2015).

Single cell analyses of muscle regeneration demonstrate the rich cellular complexity governing myogenesis and make two key observations (De Micheli et al., 2020a; Dell’Orso et al., 2019a; Giordani et al., 2019). First, in response to damage, MuSCs exit quiescence and progress through a hierarchical, dynamic myogenic program as activated MuSCs either commit to terminal differentiation or self-renew and re-acquire a quiescent state. Second, as activated MuSCs progress through myogenesis, MuSCs experience dramatic global changes in gene expression (Barruet et al., 2020). This rapid change in gene expression requires MuSCs to regulate the vast amount of newly transcribed RNA encoding fate-specifying transcription factors and skeletal muscle contractile apparatus constituents, among many other myogenic proteins.

RNA is regulated by an arsenal of abundant RNA-binding proteins (RBPs). Previous work on RBPs in myogenesis focused on specific RBP proteins, often associated with disease, and their function in regulating key myogenic transcription factors that define myogenesis (Apponi et al., 2011). However, more recently identified roles for RBPs in myogenesis include: (1) maintenance of MuSCs quiescence (Morrée et al., 2017), (2) MuSC activation and expansion (Cho and Doles, 2017; Farina et al., 2012), (3) myogenic differentiation (Hausburg et al., 2015; Vogler et al., 2018) and (4) MuSC-self-renewal (Chenette et al., 2016; Hausburg et al., 2015), demonstrating the critical role of RNA regulation throughout muscle regeneration.

One major RBP function is the regulation of RNA splicing, where RNA is targeted and bound by RBPs co-transcriptionally to guarantee correct splicing of nascent RNA transcripts (Dassi, 2017; Hentze et al., 2018). RBP-mediated splicing yields a rich diversity of RNA isotypes critical for translating proteins necessary for myogenesis (Brinegar et al., n.d.; Imbriano and Molinari, 2018; Nakka et al., 2018; Weskamp et al., 2020) The loss of specific RBPs and the resultant effects on splicing diversely affects quiescence, activation, or muscle differentiation and thus, mutations in RBPs are associated with muscular dystrophies and age-related neuromuscular diseases (Calado et al., 2000; Hinkle et al., 2019; Xue et al., 2020; Yu et al., 2009).

Amyotrophic lateral sclerosis, inclusion body myopathy, and muscular dystrophies are caused by RBP mutations leading to progressive muscle degeneration (Lukong et al., 2008; Xue et al., 2020). In these disorders, disease causing mutations frequently impair RBP splicing function (Cortese et al., 2014; Singh et al., 2018; Taylor et al., 2016) and restoring RBP splicing function prevents or delays the onset of these disorders, identifying splicing dysregulation as a key driver of pathology (Naryshkin et al., 2014; Scotti and Swanson, 2016; Wirth et al., 2020).

Despite the fact that RBPs are essential for skeletal muscle regeneration and are frequently mutated in neuromuscular disease, little is known regarding the mechanisms regulating the timing of RBP function during myogenesis or the mechanisms by which RBPs influence myogenic cell fate decisions. Using single cell RNA-sequencing we examined temporal RBP expression of several neuromuscular disease-associated RBPs in MuSCs during myogenic differentiation to clarify the timing of RPB expression during muscle regeneration and identify networks of RBPs involved in myogenic cell fate transitions.

## RESULTS

### MYOGENIC CELL FATE TRANSITIONS REVEALED BY SINGLE CELL RNA SEQUENCING

Skeletal muscle has a remarkable ability to repair following injury. In response to injury, DNA-based lineage tracing and single cell sequencing experiments demonstrate that MuSCs activate, proliferate and differentiate to produce the majority of myonuclei by 7-days post injury (dpi) (De Micheli et al., 2020). MuSCs also self-renew to replenish the quiescent MuSC stem cell population with the majority of self-renewal occurring between 5- and 8-dpi (Pawlikowski et al., 2019). Thus, between 4- and 8-dpi the MuSC population traverses all stages of myogenesis as skeletal muscle regenerates. During this time, MuSCs regulate specific networks of RNA to direct myogenic fate decisions, but the role of RBPs in maintaining a malleable myogenic transcriptome remains poorly understood. We first sought to characterize myogenesis in single cells between 4- and 7-dpi to gain insight into the timing of RBP expression during MuSC activation, proliferation, differentiation, and self-renewal.

To induce skeletal muscle regeneration, we first injured adult mouse tibialis anterior (TA) muscles with barium chloride (Caldwell et al., 1990), collected regenerating muscle at 4- dpi and 7-dpi and performed single cell RNA sequencing. Analysis of the regenerating skeletal muscle shows strong reproducibility amongst biological replicates (S1A-B) and defines the cellular composition of the regenerating skeletal muscle (Figure 1A). Following acute muscle injury, we observe the infiltration of immune cells, dynamic cycling of individual fibro/adipogenic progenitors (FAPs), and a myogenic progenitor population (Figure 1A, S1A-B). The defined cellular populations during skeletal muscle regeneration are consistent with those from published myogenic single cell sequencing datasets, which suggests that acutely injured muscle undergoes similar mechanisms of cellular regeneration irrespective of the inciting injury (Barruet et al., 2020; De Micheli et al., 2020b; Dell’Orso et al., 2019b).

**Figure 1:**
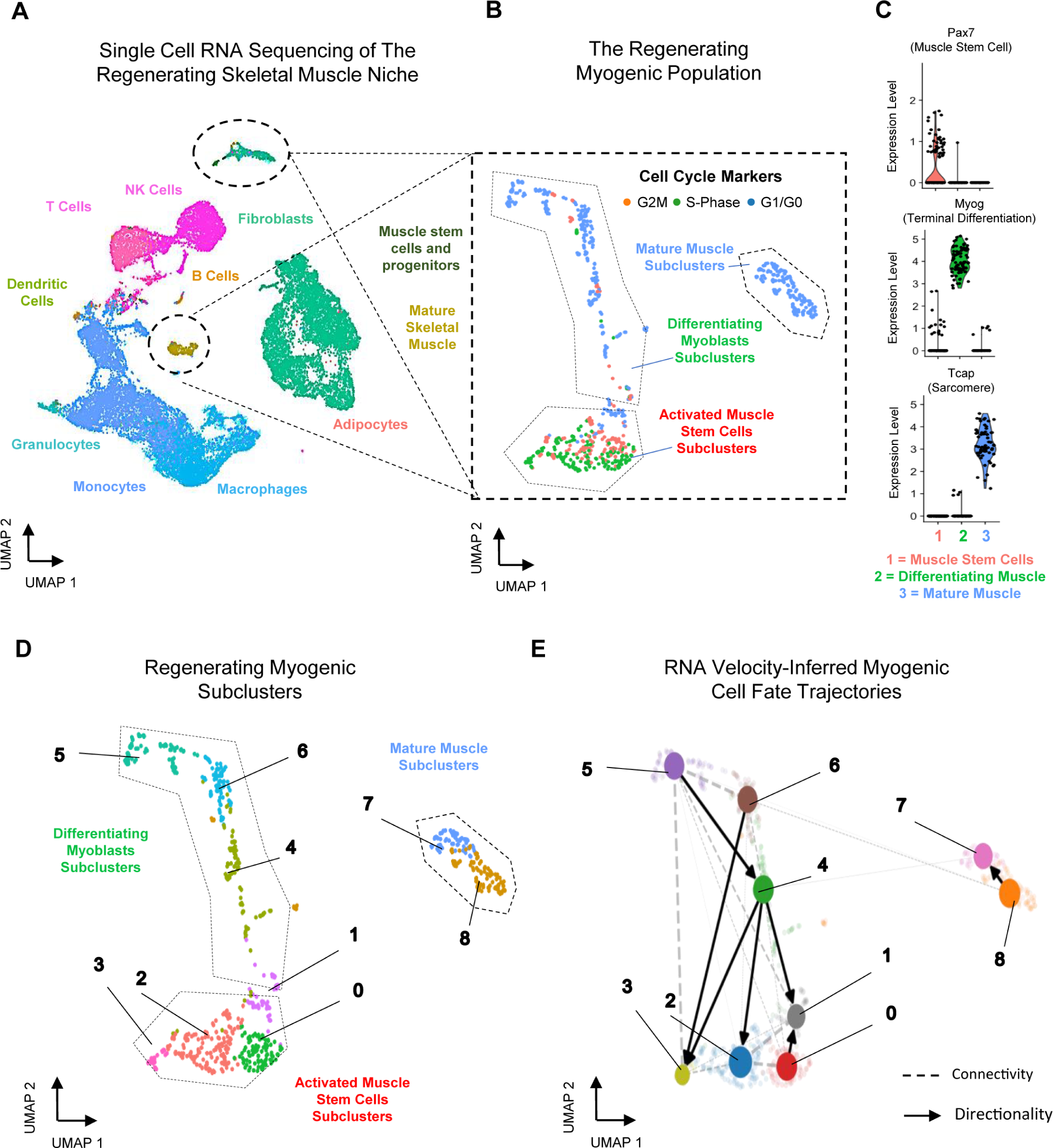
Single cell analysis reveals myogenic cell fate transitions in regenerating skeletal muscle (A) Single cell atlas of regenerating skeletal muscle at 4- and 7-days post injury (-dpi) (B) Cell cycle scoring in regenerating myogenic subclusters (C) Violin plots showing expression of myogenic markers of regeneration per myogenic clusters (D) Myogenic subclusters comprising the regenerating myogenic cellular population (E) RNA velocity-inferred myogenic cell fate trajectories. See also Figure S1.

Gene expression analysis and cell cycle scoring analysis of the myogenic populations early during muscle regeneration reveals three dominant myogenic clusters: a proliferative Pax7- positive MuSC population, a differentiating Myogenin-positive population exiting cell cycle, and a post-mitotic, terminally differentiating muscle population expressing RNAs encoding sarcomeric proteins (Figure 1B-C, S1C). These dominant myogenic populations are further subclassified into nine subclusters, which together define the regenerating myogenic population (Figure 1D). It is unclear if these sub-clusters represent specific cell state transition points within the continuum of myogenesis, or whether we are catching cells at various points along a more continuous differentiation trajectory.

Using RNA velocity, we next sought to identify temporal dynamics between each of the nine myogenic subclusters by leveraging directed dynamic information between single cells (Bergen et al., 2020). RNA velocity works on the principle that newly transcribing genes’ pre-mRNAs contain introns that have not yet be removed by RNA splicing, which can be detected in single cell data. By globally comparing pre-mRNA reads to the corresponding processed mRNA, RNA velocity can infer cellular dynamics for a single cell (La Manno et al., 2018).

We found that RNA velocity analysis on single cells identified myogenic differentiation directionality within specific subclusters, suggesting that myogenic differentiation trajectories can be resolved and predicted based on the processed and unprocessed RNA within myogenic cells (Figure S1C).

To further characterize and predict the connectivity between the nine subclusters, we utilized partition-based graph abstraction (PAGA), which estimates connectivity among different sub-clusters while preserving global data topology, inferring cellular trajectories in finer detail than traditional pseudotemporal analyses (Wolf et. al, 2019). PAGA reveals a dynamic connectivity between the myogenic subclusters 0-6 that underlies in vivo myogenesis (Figure 1E, S1D) where robust connections are observed between MuSC clusters and differentiating clusters, highlighting the underlying complexity of myogenic cell fate decisions as myogenic cells either undergo self-renewal or commit to terminal differentiation (Figure S1D). While a high degree of connectivity is observed amongst the immature myogenic populations, very few connections lead to terminally differentiated, mature muscle, suggesting that terminal differentiation may follow a singular, universal pathway rather than multiple, diverse pathways. Our analyses suggest that these subclusters likely represent dynamic cell fate transition points within proliferating MuSCs and differentiating progenitor populations, capable of either progressing to terminal differentiation or returning to a more stem-like state; however, the mechanism that drives these cell fate dynamics between myogenic subclusters remains unknown. Although the mechanism of myogenic cell fate change is clearly multifactorial, a likely driver of rapid and dynamic cell fate changes is post-transcriptional control of RNA. Our data reveal a trajectory map of early in vivo myogenesis and provide an opportunity to examine temporal RNA regulation during rapid and dynamic myogenic cell fate transitions.

### RNA-BINDING PROTEIN EXPRESSION IS TEMPORALLY DEFINED DURING MYOGENESIS

RNA is regulated in large part through the interaction with an ensemble of RBPs. While RBP expression increases during muscle regeneration, little is known about the timing of RBP expression in individual MuSCs and how specific RBP expression changes over the course of myogenesis *in vivo*. We hypothesized that temporal regulation of RBP expression in MuSCs modulates post-transcriptional gene expression to influence cell fate.

To test this hypothesis, we examined the steady state transcript levels of all RBPs in myogenic subclusters within proliferating MuSC and differentiating progenitor populations (subclusters 0-6). Hierarchical clustering of RBP transcript levels in each myogenic subcluster demonstrates that RBP expression correlates to the underlying myogenic cell state (Figure 2A). Groups of RBPs appear to exhibit similar expression patterns related to the associated myogenic subcluster (Figure 2A). Thus, coordinated upregulation or downregulation of select groups of RBPs serves to post-transcriptionally regulate RNAs specific to dynamic cell state changes over the course of myogenesis.

**Figure 2:**
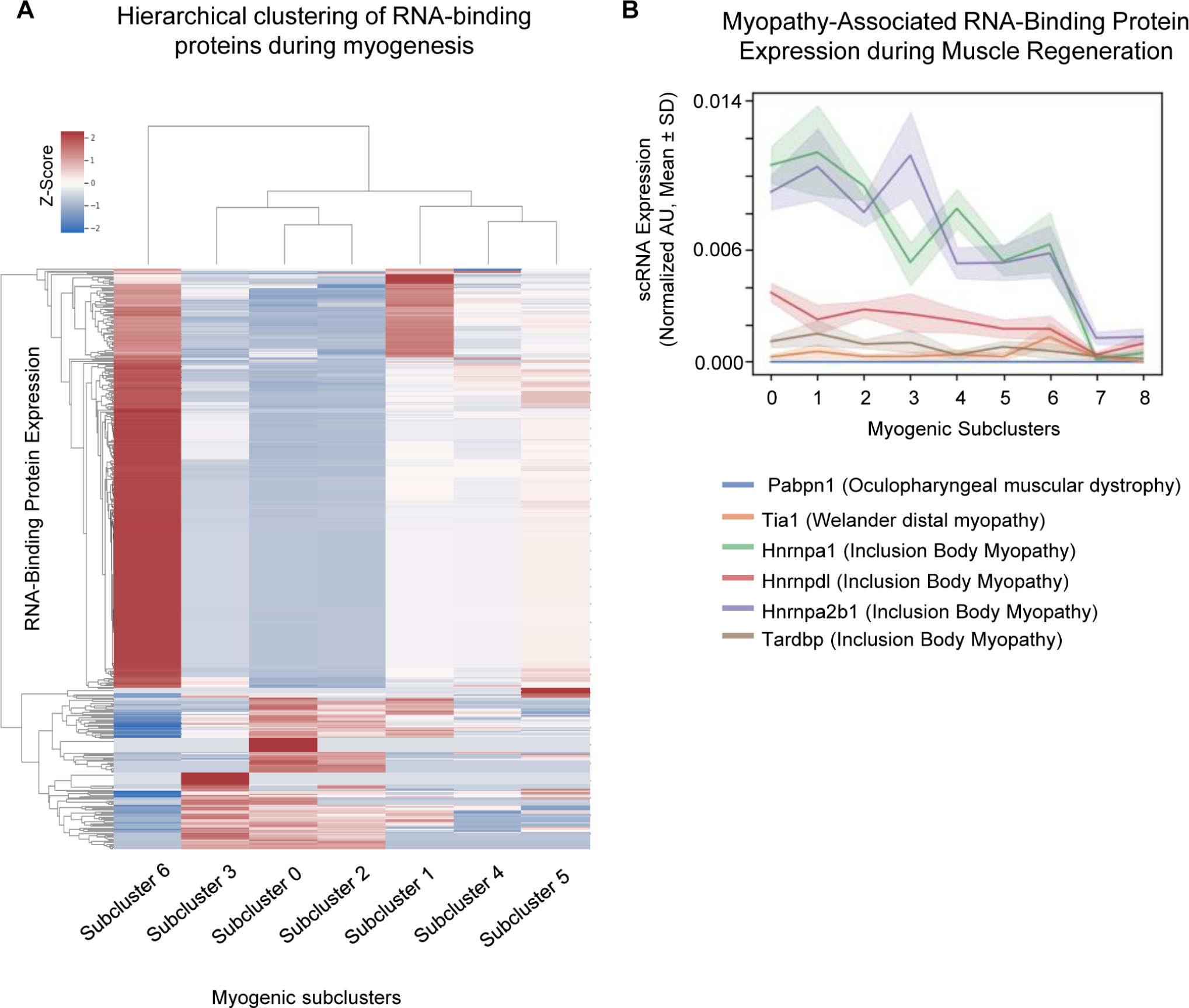
Cluster-specific RNA-binding protein expression is temporally defined during myogenesis. (A) Hierarchical clustering of RNA-binding protein expression in myogenic subclusters 1-6 during myogenesis. (B) Line plot of select myopathy-associated RNA-binding protein expression in subclusters 0-8 during myogenesis.

Dysfunction in neuromuscular disease associated RBPs gives rise to distinctive myogenic phenotypes. For example, Tardbp (TDP-43) dysfunction impairs myogenic proliferation, whereas loss of Hnrnpa1 disrupts terminal myogenic differentiation (Liu et al., 2017; Vogler et al., 2018). Since select RBP dysfunction contributes to degenerative neuromuscular diseases, defining the expression timing of disease related RBPs may help provide mechanistic clues to the contribution of RBPs to muscle pathology.

We observe cell state specific changes in the gene expression for disease associated RBPs (Figure 2B). For example, TDP-43 expression is highest in subcluster 1, Hnrnpa2b1 expression is highest in subcluster 3, and Tia1 expression is highest in subcluster 6. Expression of specific disease associated RBPs fluctuates and peaks at different time points during myogenesis, which may influence in which cell clusters RBP dysfunction is most deleterious. Intriguingly, some RBPs display marked comparative expression timing changes. For example, both Hnrnpa1 and Hnrnpa2b1 are highly expressed in subcluster 1, while in subcluster 3, Hnrnpa1 decreases and comparatively, Hnrnpa2b1 increases expression. Finally, the amount of RBP RNA expressed varies dramatically between different RBPs. For example, Hnrnpa2b1 expression is five-fold higher in proliferating MuSC and in differentiating MuSC clusters as compared to TDP-43 expression (Figure 2B). The differences in expression between subclusters may reflect functional requirements for specific RBPs at precise times during myogenic cell fate changes.

Individual RBP expression, including the expression of many RBPs linked to human disease, appears to correlate with specific myogenic cell states. Myogenic cells may alter the expression of these individual RBPs within distinct cell stages to post-transcriptionally regulate large families of RNAs. By modulating the expression of key RBPs, and in turn specific subsets of RNA, myogenic cells could more easily transition between different cell fate trajectories without massive transcriptional changes.

### HNRNPA2B1 EXPRESSION DYNAMICS IN REGENERATING SKELETAL MUSCLE

The five-fold expression increase in Hnrnpa2b1 during myoblast differentiation points towards a functional role for Hnrnpa2b1 in regulating muscle regeneration. As an RBP involved in regulating RNA splicing, stability and transport, but also found mutated in neuromuscular diseases such as inclusion body myositis, we hypothesized that Hnrnpa2b1 may function as a pivotal myogenic RNA regulator (Alarcón et al., 2015; Geissler et al., 2016; Percipalle et al., 2002). Here, Hnrnpa2b1 may influence the cycling of myogenic cells between cell fate trajectories to orchestrate successful differentiation. The function of Hnrnpa2b1 during myogenesis is unknown, but in hematopoietic stem cells, Hnrnpa2b1 regulates cell differentiation (Wang et al., 2018) and proliferation (He and Smith, 2009). The mRNA of Hnrnpa2b1 is highly expressed, and it undergoes large relative expression timing changes as compared to other RBPs between myogenic subclusters (Figure 2A-B). We therefore examined the influence of Hnrnpa2b1 on myogenic cell fate.

To validate the dynamics of Hnrnpa2b1 protein during *in vivo* skeletal muscle regeneration, mouse TA muscle was injured with barium chloride, harvested at specific time points following injury, and assayed for Hnrnpa2b1 immunoreactivity. In uninjured muscle, Hnrnpa2b1 is present in a rare subset of Pax7-postitive MuSCs and present in some peripherally located myonuclei in mature skeletal muscle fibers (Figure 3A-B, S3A). By five days post injury (5-dpi), Hnrnpa2b1 expression increases and is present in the majority of Pax7-postitive MuSCs and centrally located myonuclei of immature regenerating myofibers (Figure 3A-C, S3A-B).

**Figure 3:**
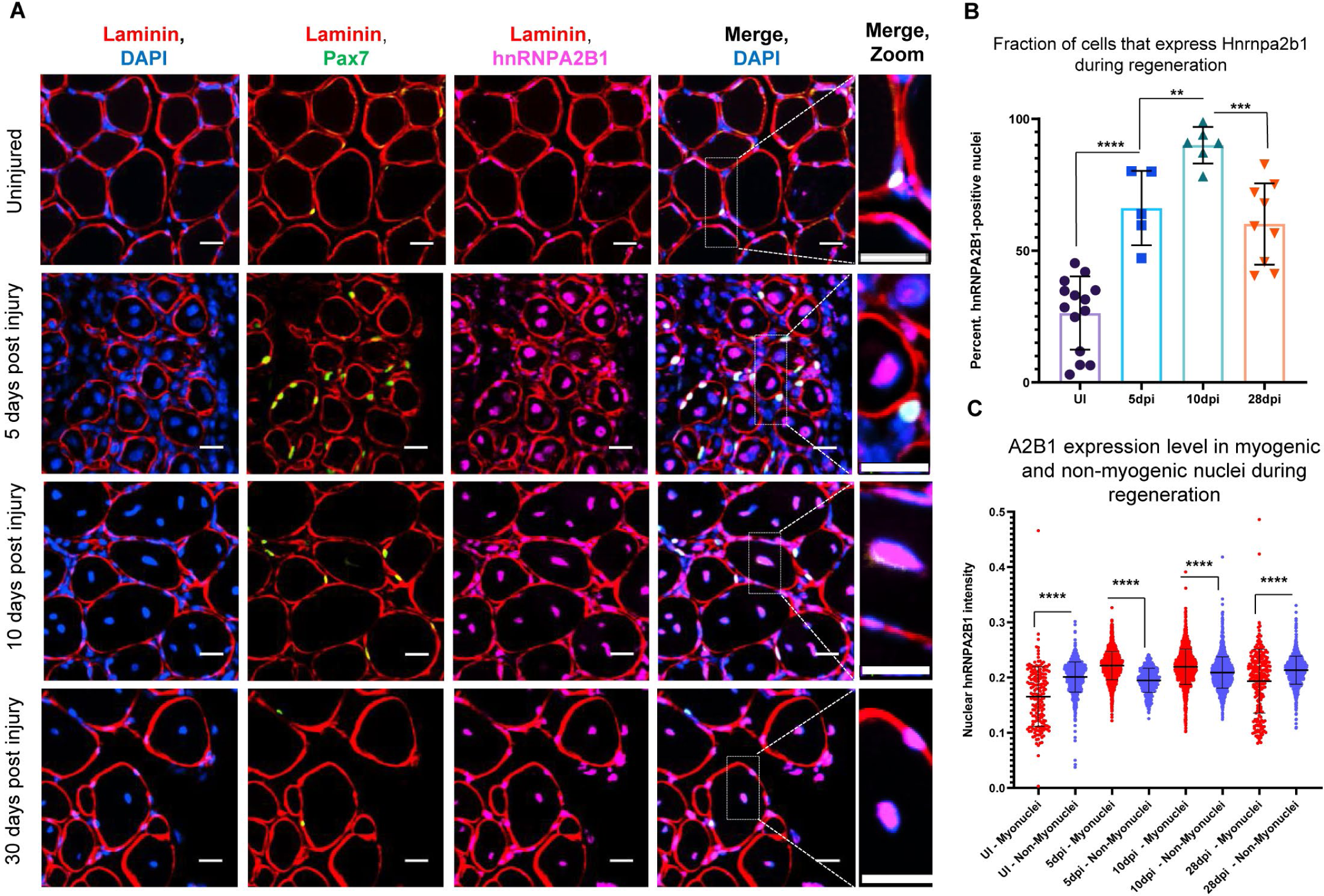
Hnrnpa2b1 is upregulated in myogenic nuclei during skeletal muscle regeneration. (A) Hnrnpa2b1 immunoreactivity in uninjured (UI), 5-,10-, and 30-dpi regenerating mouse muscle. All images represent n=3 biological replicates; scale = 20µM. (B) Nuclear Hnrnpa2b1 immunoreactivity in UI, 5-, 10-, 28-dpi regenerating muscle (C) Hnrnpa2b1 nuclear immunoreactivity intensity in either myonuclei or non-myonuclei in UI, 5-, 10-, 28-dpi regenerating muscle. See also Figure S3.

As muscle continues to regenerate, Hnrnpa2b1 levels peak in both Pax7-positive MuSCs and in regenerating myofibers at 10-dpi followed by a decline in expression as myofibers fully regenerate by 28-dpi (Figure 3A-C, S3A, S3C-D). These results identify a global increase in Hnrnpa2b1 expression early during muscle regeneration in both proliferating Pax7-positive MuSCs and in newly regenerating myofibers, correlating with *in vivo* MuSC Hnrnpa2b1 mRNA expression dynamics.

While an increase in Hnrnpa2b1 expression may be due to a global increase in RNA transcription in cells populating a regenerating muscle, Hnrnpa2b1 immunoreactivity peaks at 10-dpi with a wide, notable variation between muscle progenitor cells (Figure S3C). Thus, Hnrnpa2b1 is likely dynamically regulated during myogenesis. In comparison to Hnrnpa2b1 expression, TDP-43 immunoreactivity peaks in Pax7-positive MuSCs and regenerating myofibers at 5-dpi, before declining to near baseline by 10-dpi (Figure S3E-G). Peak TDP-43 mRNA expression occurs in a different myogenic subcluster than peak Hnrnpa2b1 mRNA expression (Figure 2B). Thus, the timing of TDP-43 mRNA and protein expression differs from that of Hnrnpa2b1 during muscle regeneration, suggesting that individual RBPs influence specific cellular aspects of *in vivo* myogenesis. Here, Hnrnpa2b1 function may be more important at later stages of myogenesis as compared to TDP-43.

### HNRNPA2B1 IS REQUIRED FOR MYOBLAST DIFFERENTIATION

The notable increases in Hnrnpa2b1 mRNA and protein expression at specific points during muscle regeneration suggest that Hnrnpa2b1 may play a functional role at specific myogenic fate transitions. To test this hypothesis, we sought to characterize the function of Hnrnpa2b1 in an *in vitro* model of myogenesis. We find Hnrnpa2b1 expressed in both proliferating myoblasts and differentiating myocytes. Analogous to Hnrnpa2b1 expression *in vivo*, Hnrnpa2b1 expression increases during myoblast differentiation, peaks by 3 days of differentiation and wanes as terminally differentiated, multinucleated myotubes continue to mature in culture (Figure 4A-B). The observation that Hnrnpa2b1 expression peaks at later stages of both *in vivo* and *in vitro* myogenesis points to a possible role for Hnrnpa2b1 in regulating differentiation.

**Figure 4:**
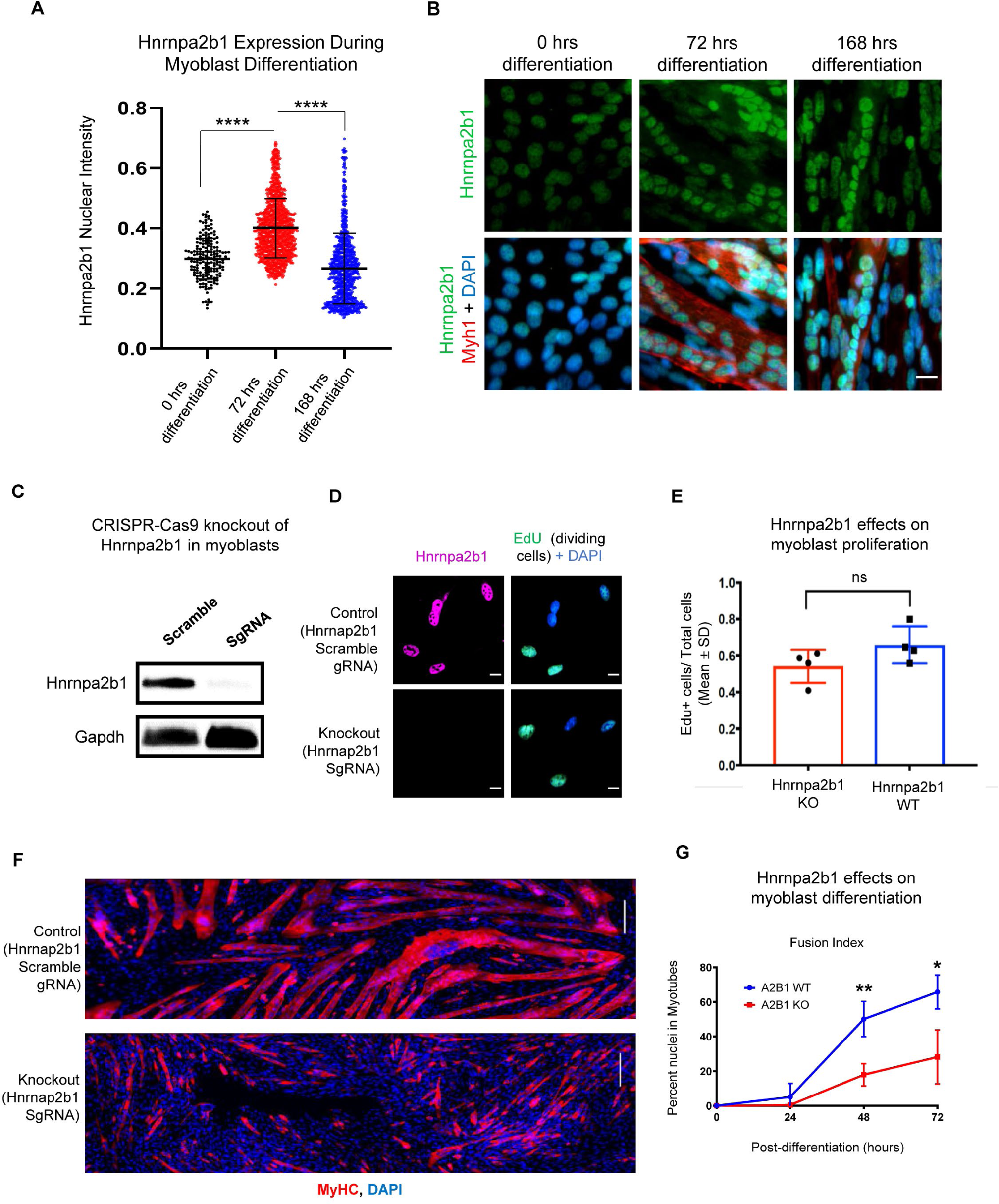
Hnrnpa2b1 is required for myogenic differentiation. (A) Hnrnpa2b1 nuclear expression in exponentially growing myoblasts (0 hours) and differentiating myotubes at 72 and 168 hours (B) Immunofluorescence showing Hnrnpa2b1 expression in myoblasts and differentiating myotubes (C) Western blot analysis of CRISPR-Cas9 knockout (KO) and scramble Hnrnpa2b1 sgRNA in C2C12 myoblasts (D) EdU-pulsed wildtype (WT) myoblasts and Hnrnpa2b1 KO myoblasts (scale = 10uM) (E) Quantification of EdU incorporation in WT and KO Hnrnpa2b1 myoblasts (ns = non-significant) (F) Staining for MyHC of differentiating in WT and KO Hnrnpa2b1 myotubes (scale = 200uM) (G) Quantification of Fusion Index (percentage of nuclei fused into myotubes) during differentiation in either WT and KO Hnrnpa2b1 myotubes. All quantified data represent mean ± SD, two-tail student t- test p-value: *=<0.05, ** =<0.01, ***=<0.001, ****=<0.0001 unless otherwise stated. See also Figure S4

To explore whether Hnrnpa2b1 functions at distinct time points during myogenesis, we knocked out Hnrnpa2b1 in myoblasts using CRISPR/Cas9 and assessed the functional consequences of Hnrnpa2b1 loss on myoblast proliferation and differentiation (Figure 4C, S4A- B). No significant difference between wild type (WT) and Hnrnpa2b1 knockout (KO) proliferating myoblasts was detected when labeled with a timed pulse of the thymidine analog 5-Ethynyl-2’-deoxyuridine (EdU; Figure 4D-E).

Loss of Hnrnpa2b1, however, impaired myoblast differentiation. After 48 hours of differentiation, wild type myoblasts had largely differentiated into multinucleated, myosin heavy chain-positive myotubes (Figure 4F-G), while Hnrnpa2b1 KO myoblasts were unable to form large multinucleated myotubes (Figure 4F-G). The Hnrnpa2b1 KO differentiation defect persisted after extended differentiation in culture, suggesting an inability to terminally differentiate (Figure 4G, S4C). Thus, Hnrnpa2b1 function is required for myotube differentiation but not myoblast proliferation, despite Hnrnpa2b1 being expressed in both differentiating myotubes and in proliferating myoblasts.

### HNRNPA2B1 IS A MYOGENIC SPLICING REGULATOR CRITICAL FOR TERMINAL MYOGENIC DIFFERENTIATION

Hnrnpa2b1 is an RNA splicing regulator and therefore, we hypothesized that Hnrnpa2b1 may function to ensure RNA splicing regulation to enable muscle differentiation. To test this hypothesis, we performed high-coverage RNA deep sequencing of Hnrnpa2b1 KO and wild type differentiating myotubes followed by differential gene expression and differential splicing analyses (Figure 5A, Supplementary Table 3 & 4). We identified 2,167 RNAs with significantly altered splicing between Hnrnpa2b1 wild type and Hnrnpa2b1 KO myotubes, of which 40% were differentially expressed, suggesting that alternative splicing may impact RNA stability of these transcripts (Figure 5A, Supplementary Table 3 & 4). These results support the hypothesis that Hnrnpa2b1 splices transcripts regulating myogenic cell fate during muscle differentiation.

**Figure 5:**
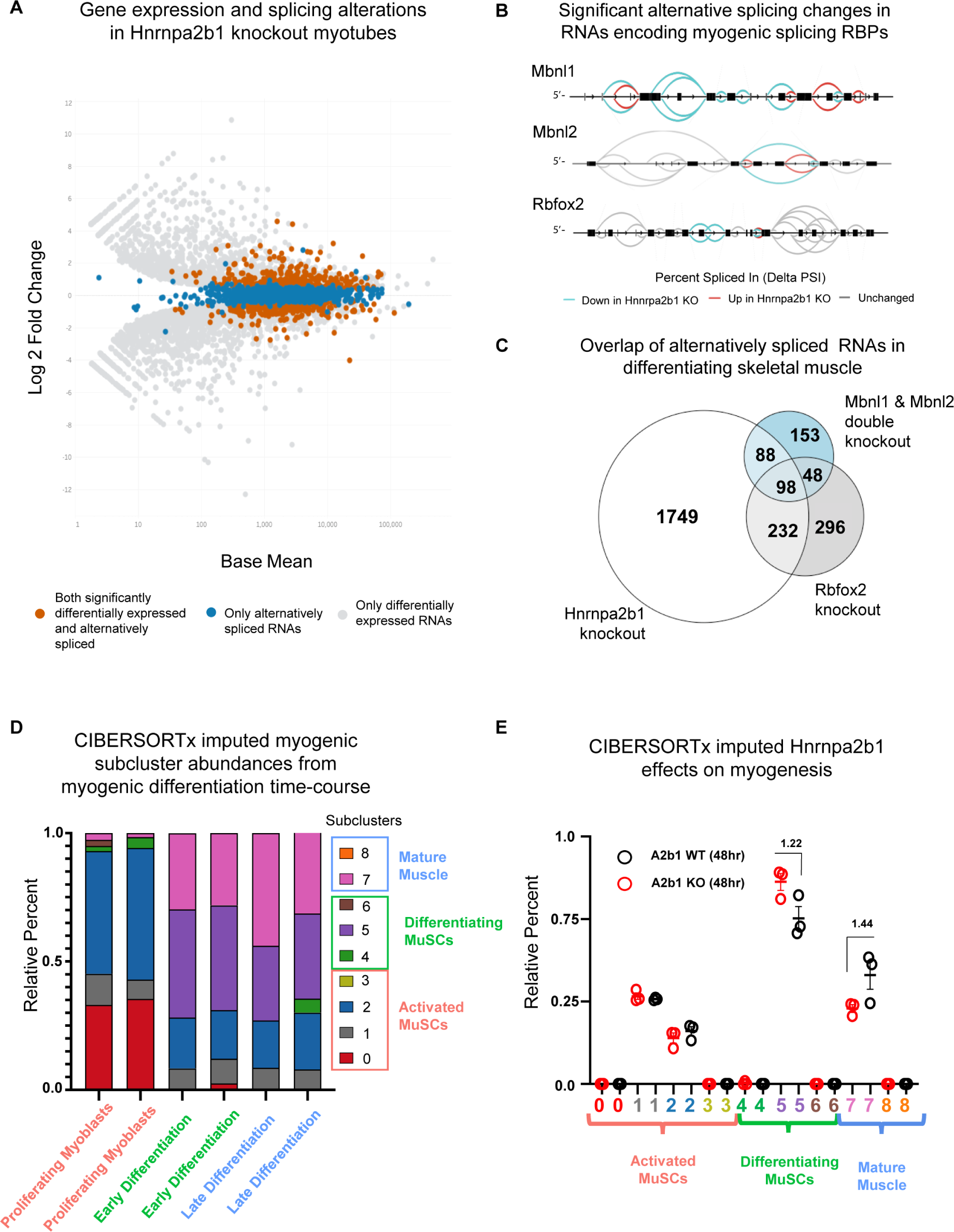
Hnrnpa2b1 is a myogenic splicing regulator critical for terminal myogenic differentiation. Differential gene expression identified by DESeq2 in WT and Hnrnpa2b1 KO myotubes with alternative splicing changes identified by Leafcutter emphasized (differential gene expression significance p- adjusted=<0.05, splicing FDR p-value =<0.05) (B) Sashimi plots of significant alternative splicing changes (dPSI) for Mbnl1, Mbnl2, Rbfox2 in differentiating Hnrnpa2b1 KO myotubes (C) Venn diagram of significantly altered spliced transcripts in Hnrnpa2b1 KO, Mbnl1/Mbnl2 double-KO (Thomas et al., 2017), and Rbfox2 KO (Singh et al., 2014) differentiating myogenic cultures (D) Myogenic-trained CIBERSORTx imputed stacked bar chart of myogenic subcluster percentages in C2C12 differentiation time course (Trapnell et al., 2010) (E) Myogenic-trained CIBERSORTx machine learning imputed percentages of myogenic clusters from WT and KO Hnrnpa2b1 differentiating myotubes. Numbers refer to fold change between myogenic cluster percentages.

These widespread splicing aberrations in Hnrnpa2b1 KO cells may also help explain their impaired differentiation.

Given the high burden of splicing alterations identified in Hnrnpa2b1 KO myotubes, and the propensity of RBPs to regulate the splicing of additional RBPs, we intuited that Hnrnpa2b1 may regulate splicing of transcripts encoding additional splicing proteins that in turn affect myogenic cell fates. Synergistic splicing regulation of related RBPs by other splicing associated RBPs cooperatively influences developmental timing and myogenesis (Klinck et al., 2014; Venables et al., 2013). Temporal regulation by splicing associated RBPs of other related RBPs may provide a mechanism to finely tune splicing dynamics and RNA processing during myogenic differentiation.

Differential splicing analysis in Hnrnpa2b1 KO reveals numerous alternatively spliced RNAs that encode splicing related proteins including the myogenic splicing regulator RBPs Mbnl1, Mbnl2, and Rbfox2 (Figure 5B). As loss of Mbnl1, Mbnl2, or Rbfox2 results in impaired myogenic differentiation, the alternative splicing of these proteins may compromise their splicing function. Consistent with this prediction, we observe a significant overlap in the alternatively spliced RNA populations identified between Hnrnpa2b1 and Mbnl1 & Mbnl2 double knockout differentiating muscle (hypergeometric p-value = 1.2x10^-77^), and Hnrnpa2b1 and Rbfox2 knockout differentiating muscle (hypergeometric p-value = 6.1x10^-143^) (Figure 5C). Further, comparing the precise splice site and effect of the splicing alteration (inclusion or exclusion of introns) in shared alternatively spliced RNAs identifies many co-regulated splicing changes shared between Mbnl1, Mbnl2, Rbfox2, and Hnrnpa2b1. These RBPs co-regulate the splicing in –cis of shared RNA targets (Figure S5A), implying a role for Hnrnpa2b1 as a central regulator within a network of splicing factors that directs specific myogenic cell fate decisions by synergistically regulating a common cadre of target RNAs.

The potential redundancy between RBPs and their splicing functions is consistent with our earlier observations showing aligned expression patterns for related RBPs which correlate with the underlying myogenic subcluster (Figure 2A). As these RBPs are differentially expressed in distinct subclusters corresponding to the stage of myogenic differentiation, these RBPs may regulate the transition from one subcluster to the next. Therefore, we predict that defects in a given RBP may disrupt normal myogenesis and result in myogenic cells arresting in distinct subclusters during differentiation. To test this, we characterized the impact of Hnrnpa2b1 knockout on myogenic subcluster abundances during differentiation, where we employed the machine learning-based methodology CIBERSORTx to computationally impute the abundance of myogenic subclusters present in differentiating Hnrnpa2b1 KO and WT cells. CIBERSORTx estimates the abundance of cell types in a mixed cell population from bulk RNA sequencing using single cell RNA sequencing data as a training set (Newman et al., 2019). By training this machine learning classifier on our single cell myogenic data, CIBERSORTx could calculate the percentages of cells at different stages of myogenesis from bulk RNA sequencing. If a block in differentiation arises due to RBP dysfunction or knockout, cells would be expected to accumulate into subclusters behind this block. Myogenic-trained CIBERSORTx would then predict the increase in the percentages of these cellular subclusters, pinpointing the timing of differentiation defects due to loss of RBP function.

As a proof-of-concept, we trained CIBERSORTx on our myogenic single cell dataset and imputed the percentages of myogenic cells from an RNA-sequencing dataset of C2C12 cell differentiation performed on exponentially growing myoblasts, and differentiating myotubes at 60 hours and 120 hours after differentiation induction (Trapnell et al., 2010). If myogenic-trained CIBERSORTx is capable of classifying distinct states of myogenic differentiation represented in MuSC single cell sequencing data from bulk C2C12 cell RNA sequencing, then proliferating MuSCs should cluster with proliferating C2C12 cells and MuSCs undergoing differentiation are expected to cluster with differentiating C2C12 cells, proportional to the time of differentiation the RNA sequencing was performed. Indeed, myogenic-trained CIBERSORTx reveals an enrichment for proliferative MuSCs subclusters in the exponentially growing C2C12 myoblast population and a shift towards differentiating subclusters proportional to differentiation duration (Figure 5D, S5B, Supplementary Table 5). The observation that not all myogenic subclusters were identified by CIBERSORTx in the C2C12 differentiation time course (e.g. subcluster 3) suggests that some subclusters are unique to myogenesis *in vivo* or do not comprise a substantial fraction of the *in vitro* C2C12 population at the assessed time points (Figure 5D, S5B).

Myogenic-trained CIBERSORTx detects a small percentage of differentiated cells (e.g. subcluster 7) present in the exponentially growing myoblasts (Figure 5E, S5B-C). Consistent with this observation, we performed immunohistochemistry against MyoG, a marker of differentiation, in exponentially proliferating myoblasts and identify a similar small percentage of MyoG expressing cells in exponentially growing myoblasts as predicted by CIBERSORTx (Figure S5C). These results demonstrate that myogenic-trained CIBERSORTx is capable of accurately imputing myogenic subcluster abundances from bulk RNA sequencing data.

Next, we sought to test the impact of Hnrnpa2b1 loss on cell subclusters during myogenesis. Myogenic-trained CIBERSORTx identifies an enrichment for subcluster 5 myogenic cell signatures and a concomitant decrease in terminally differentiated subcluster 7 in Hnrnpa2b1 KO cells (Figure 5E) showing that Hnrnpa2b1 splicing function ensures MuSCs transition from subcluster 5 to subcluster 6 during terminal differentiation.

### RBP ENGAGEMENT SCORING DELINEATES FUNCTIONAL TIMING OF RBPS DURING MYOGENESIS

The timeframe in which a splicing RBP performs its function is tied to both the expression timing of the RBP and the expression of the RBP RNA targets. Hnrnpa2b1 promoting the transition of MuSCs from subcluster 5 to subcluster 6 during terminal differentiation suggests that RBP function may be linked to a particular cell state. The necessity of Hnrnpa2b1 for terminal differentiation may reflect both the specific temporal expression of Hnrnpa2b1 as well as Hnrnpa2b1 target RNAs. Therefore, we next identified and defined the expression dynamics of Hnrnpa2b1 RNA targets during myogenesis.

We performed Hnrnpa2b1 enhanced UV crosslinking and immunoprecipitation (eCLIP) in proliferating myoblasts and in differentiating myotubes (Figure 6A, S6A-C) to identify Hnrnpa2b1-associated RNAs. Hnrnpa2b1 eCLIP reveals 808 binding sites across 247 genes for myoblasts and 1,030 binding sites across 137 genes for myotubes, which are significantly enriched compared to size-matched input (reflecting all RNA–protein interactions in the input) (Figure S6D, Supplementary Table 2). Hnrnpa2b1 RNA target eCLIP peaks were highly correlated between biological replicates, had thousands of reproducible eCLIP clusters by irreproducible discovery rate analysis, and capture prior identified Hnrnpa2b1 RNA targets including Hnrnpa2b1’s own 3’ UTR (Figure S6D-G) (Martinez et al., 2016). Analysis of Hnrnpa2b1 RNA binding sites reveals an enrichment for binding sites in the 3’UTR and proximal introns of target RNAs (Figure 6A). Using native Hnrnpa2b1 RNA immunoprecipitation (RIP) followed by RT-qPCR of eCLIP target RNAs, we validated the interaction of Hnrnpa2b1 with several of Hnrnpa2b1 target RNAs identified by eCLIP (Figure 6B, S6H, Primers used detailed in Supplementary Table 1).

**Figure 6:**
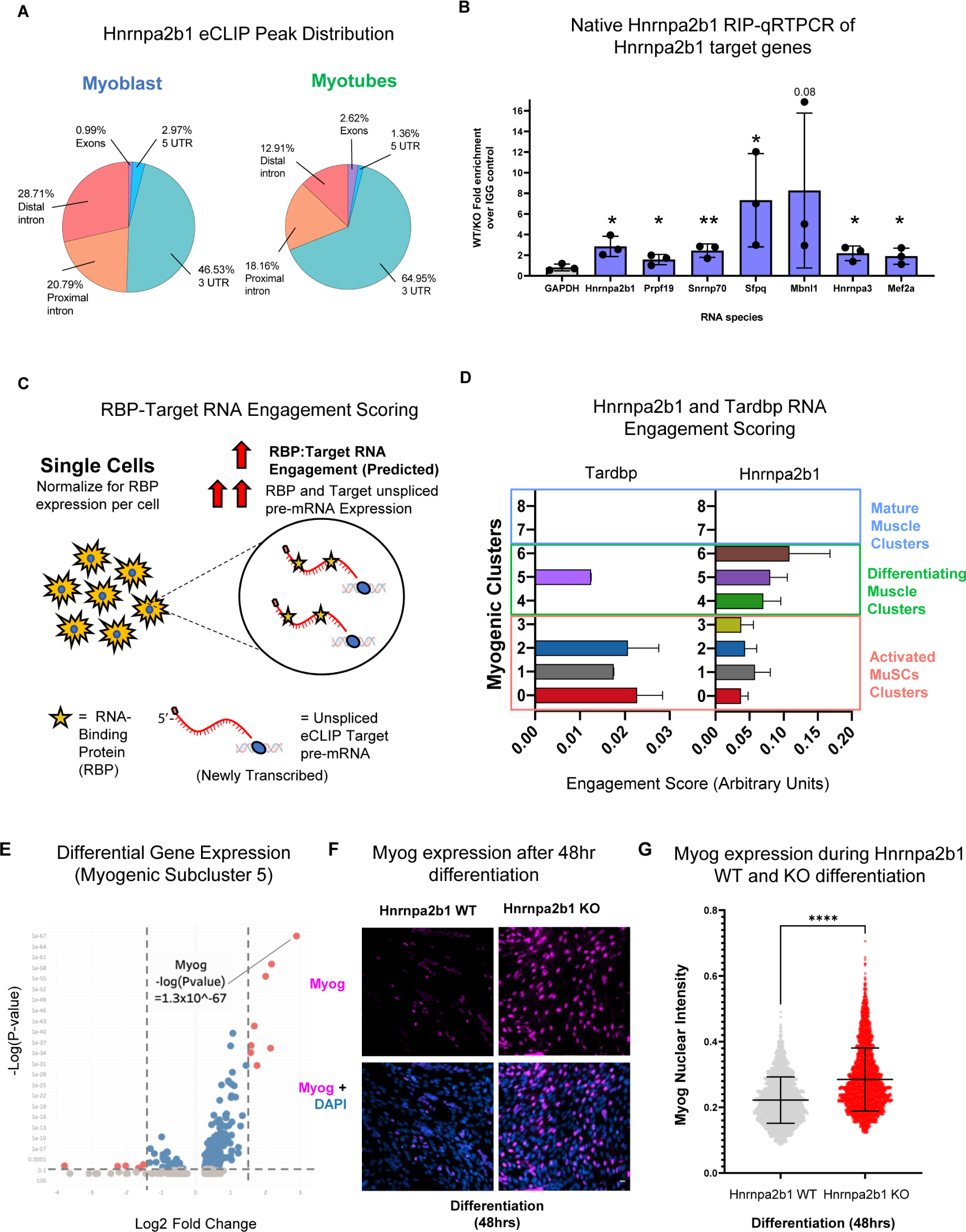
RNA engagement scoring delineates functional timing of RBPs during myogenesis. (A) Genomic distribution of Hnrnpa2b1 eCLIP peaks in myoblasts and myotubes (B) Native Hnrnpa2b1RNA immunoprecipitation (RIP)-qRT-PCR of RNA targets identified by eCLIP (C) Schematic for RBP-Target RNA engagement scoring (D) RNA-binding protein (RBP) – Target RNA engagement scoring of Tardbp and Hnrnpa2b1in myogenic subclusters (median ± 95% CI for across all RBP-RNA engagement scores in each myogenic subcluster) (E) Differential gene expression analysis of myogenic subcluster 5 (F) Myog staining in WT and Hnrnpa2b1 KO 48 hour differentiated myotubes (G) Myog intensity in differentiating WT and Hnrnpa2b1 KO myotubes (mean ± SD, two-tailed student t-test p-value: ****=<0.0001). See also Figure S6.

Many RBPs with splicing function, including Hnrnpa2b1, engage with target RNAs co-transcriptionally to regulate RNA splicing (Fu and Ares, 2014). Since recent computational methods allow for newly transcribed RNA or pre-mRNA expression to be measured at a single cell level, we may be able to infer the degree of RBP functionality within an individual cell by correlating the expression of a given RBP to the expression of the newly transcribed RBP target pre-mRNAs. We termed this approach RBP engagement scoring (Figure 6C).

To test if RBP engagement scoring can infer known RBP phenotypes, we first calculated RBP engagement scores for TDP-43, a well characterized splicing RBP that is essential for MuSC proliferation (Vogler et al., 2018). TDP-43 engagement scoring should be higher in proliferating MuSC clusters as compared to differentiating MuSC clusters. The pre-mRNA expression of TDP-43 target RNAs previously identified by eCLIP was quantified across all myogenic cells identified in our single cell sequencing dataset (Vogler et al., 2018). TDP-43 target pre-mRNA expression was then correlated with TDP-43 expression per cell to assign each cell a TDP-43 engagement score. Next, TDP-43 engagement scores were summed across each cell within a myogenic subcluster to determine the relative activity of TDP-43 at each stage of myogenesis. We observe greater TDP-43 engagement scoring in proliferating MuSC clusters as compared to differentiating clusters or mature muscle clusters, consistent with a requirement of TDP-43 for MuSC proliferation (Figure 6D).

We next sought to use RBP engagement scoring to investigate the regeneration stage-specific timing of Hnrnpa2b1 function during myogenesis. In contrast to TDP-43, Hnrnpa2b1 engagement scoring revealed increased engagement scores in differentiating myogenic clusters (e.g. subcluster 6) and to a lesser extent in proliferating MuSCs (e.g. subcluster 1) as compared to mature muscle (Figure 6D). Hnrnpa2b1 expression and Hnrnpa2b1 target RNA expression is higher in specific differentiating subclusters, suggesting that Hnrnpa2b1 interaction with target RNAs may be increased in those subclusters. Consistent with our prior myogenic CIBERSORTx analysis, Hnrnpa2b1 activity may be increased in select myogenic subclusters during differentiation, which may explain why the loss of Hnrnpa2b1 impairs MuSC transitions from one subcluster to the next (Figure 4D-G). As Hnrnpa2b1 engagement scores are increased in differentiating myogenic subclusters (e.g. subcluster 6), loss of Hnrnpa2b1 may impair these myogenic cells’ progress towards mature muscle, leading to a build-up of differentiated but immature myogenic cells. Hnrnpa2b1 engagement scoring predicts that Hnrnpa2b1 targets are most prevalent in myogenic cluster 6 and therefore, cells may accumulate in preceding subcluster 5 in the absence of Hnrnpa2b1.

To identify gene expression signatures unique to each cluster, we performed differential gene expression analysis on myogenic subclusters 5 and 6. The myogenic transcription factor, Myogenin (MyoG) (p-value = 1x10-67) is significantly enriched in subcluster 5, which is consistent with subcluster 5 representing a key branch point between terminal differentiation and self-renewal (Figure 6E, S6I). When differentiated Hnrnpa2b1 KO and WT myoblasts were differentiated and assessed for MyoG immunoreactivity, MyoG-positive cells were increased in Hnrnpa2b1 KO cells compared WT cells, consistent with accumulation of Hnrnpa2b1 KO cells in subcluster 5 that are unable to progress or delayed in progressing to subcluster 6 (Figure 6F- G). Thus, RBP engagement scoring identifies the precise myogenic subcluster in which Hnrnpa2b1 is most functional and provides insight into how Hnrnpa2b1 loss can disrupt myogenic fate decisions.

## DISCUSSION

RBPs are essential for skeletal muscle regeneration and RBP dysfunction is associated with neuromuscular disease (Apponi et al., 2011; Farina et al., 2012). While select RBPs play an important role in myogenesis, the exact differentiation stage-specific roles of RBPs remain poorly understood.

Single cell RNA sequencing of regenerating MuSC population points to MuSCs progressing through a dynamic program to direct cell fates either towards self-renewal or terminal differentiation. To our surprise, global and neuromuscular disease associated RBP expression profiles correlate with the underlying MuSC cell state. Utilizing a combination of machine learning and novel RBP engagement scoring, we predict and experimentally validate the role of Hnrnpa2b1, a neuromuscular disease associated RBP, in orchestrating terminal myogenic differentiation. Our results suggest that RBP splicing proteins coordinately guide myogenic cell fate decisions during muscle regeneration. The coordinated expression of related RBPs may provide an additional layer of post-transcriptional gene expression regulation, which may also occur in other developmental or tissue regenerative contexts.

We propose a model in which temporally defined RBP function is influenced both by the temporal expression of an RBP as well as the temporal expression of RBP RNA targets (Figure 7). In this model, RBP regulation is achieved by the sum of RNA targets available for a given RBP to interact with. During muscle regeneration, MuSCs rapidly respond to differentiation cues. Here, MuSCs may preemptively express cell-stage appropriate RBPs to rapidly regulate a dynamic myogenic transcriptome. We liken this mechanism to the pausing of transcription factors or polymerases at the sites of transcription awaiting the right cellular cues. By having RBPs primed to function, resident MuSCs are capable of more rapidly and flexibly regulating cell fate without the need to globally alter the underlying transcriptome.

**Figure 7:**
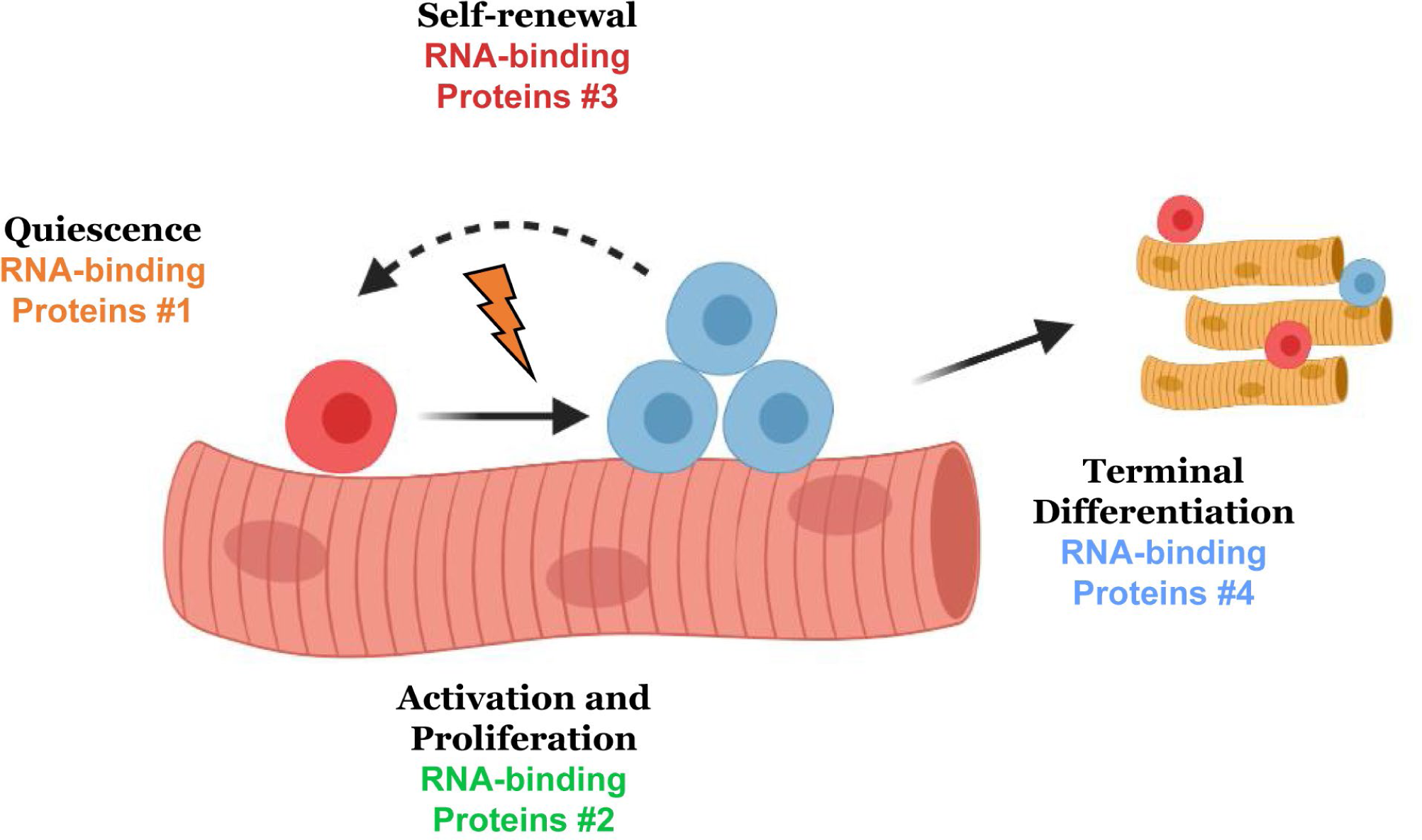
Discrete RNA-binding protein functional timing and expression as a dynamic post- transcriptional regulatory mechanism for directing myogenic fate decisions.

Our results further suggest that the function of Hnrnpa2b1 in regulating terminal differentiation may be tied to the splicing regulation of RNAs encoding additional splicing regulatory RBPs. As RBP expression is temporally defined during myogenesis, the regulation of RBP splicing by related RBPs may restrict RBP function to when only absolutely necessary.

Finally, the observation that RBP expression is temporally restricted during myogenesis implies that distinct classes of RBPs may be critical to promote specific cell fate changes during myogenesis. Similar to the stepwise waves of myogenic transcription factor expression, RBP expression waves may globally regulate post-transcriptional gene expression. Further, splicing regulation of RBPs by other RBPs may enable isoform specific RBP functions, potentially to more finely tune the myogenic transcriptome throughout differentiation. Our findings help to resolve the seemingly disparate impact that different RBPs have on myogenesis. As RBP functions may be temporally restricted, dysregulation of an individual RBP may impair myogenesis at a precise cell fate transition. Thus, temporally defined and muscle regeneration stage-specific RBP function may provide ample opportunities for therapeutic intervention in neuromuscular disease and may instruct stem cell fate decisions during tissue regeneration (Ferlini et al., 2021).

## Acknowledgments

We thank J. Dragavon, J. Orth for help with microscopy and the CU Microscopy Cores (supported by NIST-CU 70NANB15H226, NIH1S10RR026680-01A1) as well as the University of Colorado Cancer Center Genomics Core (supported by NIH- P30CA46934). We thank Theodore E. Ewachiw and Yang Zhao for their technical assistance.

## Author contributions

J.R.W, O.N.W., T.O.V., B.B.O. and R.P. conceived and designed the research, wrote the manuscript and all authors edited drafts. J.R.W., O.N.W, B.P., N.D.B., T.E. A.C., R.O., and K.J. performed 10X sc-RNA-seq, and analyzed mouse regeneration and myotube formation experiments. E.N. and J.R.W. performed eCLIP and subsequent analysis. A.T., K.R.P., K.E.Y., E.L. provided analytical support for data analysis. T.R., H.Y.C., and A.J. provided scientific insights and materials.

## Data and materials availability

All data is available in the main text or the supplementary materials. Sequencing data deposited as follows: 10X scRNA sequencing (awaiting accession number), Hnrnpa2b1 eCLIP (GSE106553), Hnrnpa2b1 WT and KO RNA sequencing (GSE152467).

## SUPPLEMENTARY MATERIALS

### MATERIALS AND METHODS

#### Mice

Mice were bred and housed according to the National Institutes of Health guidelines for the ethical treatment of animals, in a pathogen free facility at the University of Colorado at Boulder. Wild-type mice were genotype C57BL/6 (Jackson Laboratories). Cells and tibialis anterior (TA) muscles were isolated from 3-6-month-old male and female wild-type mice.

#### Mouse injuries

Male and female mice between 3-6 months of age were anesthetized with isoflurane and the left TA muscle was injected with 50μL of 1.2% BaCl2. The injured and contralateral TA muscles were harvested at the indicated time points.

#### TA collections and cell isolations

TA muscles were dissected and placed into 400U/mL collagenase at 37°C for 1h with shaking and then placed into Ham’s F-12C supplemented with 15% horse serum to inactivate the collagenase. Cells were passed through three strainers of 100µm, 70µm, and 40µm (BD Falcon) and flow through was centrifuged at 1500×g for 5 min and the cell pellets were re-suspended in Ham’s F-12C. To remove dead cells and debris, cells were passed over the Miltenyi, dead cell removal kit columns (Cat# 130-090-101). To remove RBCs, cells were incubated with antiTer119 micro magnetic beads and passed over a Miltenyi column (Cat#130-049-901). For the adult and aged uninjured TAs 6 TA muscles (from 3 mice) were pooled together. For the injured TA muscles 2 TA muscles from 2 different mice were pooled together. Cells were then counted using a BioRad TC20 automated cell counter and processed with the 10X genomics single cell sequencing kit.

#### Single Cell Sequencing

To capture, label, and generate transcriptome libraries of individual cells we used the 10X genomics Chromium Single Cell 3’ Library and Gel Bead Kit v2 (Cat#PN-120237) following the manufactures protocols. Briefly, the single cell suspension, RT PCR master mix, gel beads, and partitioning oil were loaded into a Single Cell A Chip 10 genomics chip, placed into the chromium controller, and the chromium Single Cell A program was run to generate GEMs (Gel Bead-In-EMulsion) that contain RT-PCR enzymes, cell lysates and primers for illumine sequencing, barcoding, and poly-DT sequences. GEMs are then transferred to PCR tubes and the RT-PCR reaction is run to generate barcoded single cell identified cDNA. Barcoded cDNA is used to make sequencing libraries for analysis with Illumina sequencing. We captured 1709 cells from young uninjured, 3459 from the aged uninjured, 5077 from the 4-d post injury and 2668 from the 7-d post injury. Sequencing was completed on an Illumina NovaSeq 6000, using paired end 150 cycle 2x150 reads by the genomics and microarray core at the University of Colorado Anschutz Medical Campus.

#### Single Cell Informatics

Preprocessing was performed using Cellranger v3.0.1 (10X Genomics) count module was used for alignment using cellranger-mm10-3.0.0 refdata, filtering, barcode counting and UMI counting of the single cell FASTQs. Postprocessing was performed using Seurat according to standardized workflows (Butler et al., 2018; Stuart et al., 2019). In brief, using RStudio (v4.0.0), Seurat objects were created for each Cellranger processed sample by importing filtered_gene_bc_matrices. Multiple Seurat objects then were merged, filtered, normalized, feature selected, scaled, and clustered. Non-linear dimensional reduction was performed using UMAP.

**RNA velocity** using the scVelo pipeline was performed according to standardized workflows (La Manno et al., 2018). In brief, Velocyto was run in Python v3.6.3 using Samtools v1.8, cellranger- mm10-3.0.0 refdata, and masking repetitive elements to generate unspliced/ spliced/ ambiguous read count Loom file for each 10X cellranger preprocessed library. Seurat objects of myogenic clusters (.rds files) were converted in RStudio to loom files using LoomR. In a virtual environment, Velocyto Loom files were concatenated and merged with Seurat exported myogenic Loom in Scanpy. The scVelo pipeline was then used to perform log normalization and filtering. RNA velocity was then imputed using stochastic modeling. RNA velocities were projected on pre-computed UMAP embeddings and annotated clusters.

**RBP Engagement Scoring** was performed by first determining the raw unspliced reads for every gene at single cell level using Veloctyo as above. Metadata containing raw reads counts and unspliced reads imputed using Veloctyo were exported from Scanpy anndata objects for single cells contained within myogenic clusters were exported and reimported into RStudio (v.4.0.0). To calculate an RBP-RNA engagement score for a given RBP, CLIP target genes of a given RBP genes were subset and unspliced read counts and total RBP counts were normalized to total read counts in a given cell. Scoring then represents the fraction of normalized unspliced reads of target RNAs to normalized amounts of a given RBP. To extrapolate to a given cluster, scores were then summed across all cells in a given cluster and data was represented by the median of these scores for a given cluster of cells. Target genes were identified using publicly available eCLIP datasets for TDP-43.

**CIBERSORTx** of myogenic clusters was performed according to published workflows (Newman et al., 2019). In brief, a single cell reference txt file was created from our Seurat processed single cell RNA-seq dataset. A mixture file for bulk RNA sequencing samples was created using TPM values extracted from StringTie serve as input. A single cell reference txt matrix of cells organized by myogenic clusters was then used to train CIBERSORTx using default parameters (Table S3). Cell fractions were then imputed using 100 permutations for significance analysis.

#### Immunofluorescence staining of tissue sections

TA muscles were dissected, fixed on ice for 2h with 4% paraformaldehyde, and then transferred to PBS with 30% sucrose at 4°C overnight. Muscle was mounted in O.C.T. (Tissue-Tek®) and cryo-sectioned on a Leica cryostat to generate 10μm thick sections. Frozen tissues and sections were stored at -80°C until immunofluorescent staining. Prior to immunofluorescent staining, tissue sections were post-fixed in 4% paraformaldehyde for 10 minutes at room temperature (RT) and washed three times for 5 min in PBS. Staining with the anti-Pax7, Laminin and Hnrnpa2b1 antibodies required heat-induced epitope retrieval, which was performed by placing post-fixed slides in citrate buffer, pH 6.0, and subjected them 6 min of high pressure-cooking in a Cuisinart model CPC-600 pressure cooker. For immunostaining, tissue sections were permeabilized with 0.25% Triton-X100 (Sigma) in PBS containing 2% bovine serum albumin (Sigma) for 60 min at RT. Incubation with primary antibody occurred at 4°C overnight followed by incubation with secondary antibody at room temperature for 1hr. Primary antibodies included anti-Pax7 (Developmental Studies Hybridoma Bank, University of Iowa, USA) at 1:750, rabbit anti-laminin (Sigma-Aldrich) at 1:200 and a mouse anti-Hnrnpa2b1 (ab6102, Abcam) at 1:200. Alexa-fluor secondary antibodies (Molecular Probes) were used at a 1:1000 dilution. For analysis that included EdU detection, EdU staining was completed prior to antibody staining using the Click-iT EdU Alexa fluor 488 detection kit (Molecular Probes) following manufacturer protocols. Sections were incubated with 1 μg/mL DAPI for 10 min at room temperature then mounted in Mowiol supplemented with DABCO (Sigma-Aldrich) or ProLong Gold (Thermo) as an anti- fade agent.

#### Cell culture and growth conditions

C2C12 myoblast cells: Immortalized murine myoblasts (American Type Culture Collection) were maintained on uncoated standard tissue culture plastic or gelatin-coated coverslips for imaging experiments at 37°C with 5% CO2 in DMEM with 20% fetal bovine serum and 1% penicillin/streptavidin. To induce myoblast differentiation and fusion into myotubes, C2C12 myoblasts at 80% confluence were switched to DMEM media supplemented with 5% horse serum, 1% penicillin/streptavidin and 1X Insulin-Transferrin-Selenium in DMEM.

#### Hnrnpa2b1 CRISPR-Cas9 knockout and EdU incorporation

CRISPR-Cas9 knockout was performed in C2C12 myoblasts. Single guide RNA (sgRNA) against Hnrnpa2b1 (5’- GAGTCCGCGATGGAG) were designed using (crispr.mit.edu) and cloned into pSpCas9(BB)- 2A-Puro (PX459). C2C12 myoblasts were transfected with JetPrime using standard protocols. Myoblasts that integrated the CRISPR construct were selected with puromyocin (1 μg/mL) for one week. Myoblasts KO and WT clones were isolated using cloning ring and selectively detaching clonal populations via trypsinization. Clonal population KO was validated via immunofluorescence and western blotting against Hnrnpa2b1 with anti-Hnrnpa2b1 antibodies (ab6102 and ab31645). EdU incorporation: C2C12 myoblasts were incubated with 10uM 5- ethynyl-2’-deoxyuridine (EdU – Life Technologies) for three hours. Cells were washed, fixed, and stained using the methods described below.

#### Immunofluorescence staining of cells

C2C12 myoblast cells were washed with PBS in a laminar flow hood and fixed with 4% Paraformaldehyde for 10 min at room temperature in a chemical hood. Cells were permeabilized with 0.25% Triton-X100 in PBS containing 2% bovine serum albumin (Sigma) for 1 hour at RT. Cells were incubated with primary antibody at 4°C overnight, then incubated with secondary antibody at room temperature for 1hr. Primary antibodies included mouse anti-Hnrnpa2b1 (ab6102, Abcam) at 1:200, mouse-anti Myogenin (ab82843, Abcam) and a mouse anti-MHC MF-20 (Developmental Studies Hybridoma Bank, University of Iowa, USA) at undiluted, neat concentration. Alexa fluor secondary antibodies (Molecular Probes) were used at a 1:1000 dilution.

#### Microscopy and image analyses

Images were captured on a Nikon inverted spinning disk confocal microscope. Objectives used on the Nikon were: 10x/0.45NA Plan Apo, 20x/0.75NA Plan Apo and 40x/0.95 Plan Apo. Confocal stacks were projected as maximum intensity images for each channel and merged into a single image. Brightness and contrast were adjusted for the entire image as necessary against secondary antibody treated control immunofluorescent sections. Cellprofiler was used to quantify IHC and IF images using custom analysis pipelines unless otherwise stated.

#### Western blotting of cell and tissue lysates

Western blot was performed according to standard protocols. Equal volumes (20 µl) of fractions were then resolved on a 4–12% Bis-Tris SDS– PAGE gel and transferred to a nitrocellulose membrane (Bio-Rad). Primary antibodies included mouse anti-Hnrnpa2b1 (1:200; ab6102, Abcam) and Monoclonal rabbit anti-GAPDH (14C10) conjugated to HRP (1:2000; Cell Signaling, 3683S).

#### Hnrnpa2b1 eCLIP sequencing

C2C12 myoblasts were seeded at 6 x 10^6^ cells per 15 cm plate, grown 24hrs at 37C, 5% CO2 and either harvested (undifferentiated myoblasts) or differentiated in differentiation media for 4 days. Hnrnpa2b1 enhanced CLIP (eCLIP) was performed according to established protocols.(Nguyen et al., 2018; Van Nostrand et al., 2016) In brief, Hnrnpa2b1-RNA interactions were stabilized with UV crosslinking (254 nm, 150mJ/cm^2^). Cell pellets were collected and snap frozen in liquid N2. Cells were thawed, lysed in eCLIP lysis buffer (50 mM Tris-HCl pH 7.4, 100 mM NaCl, 1% NP-40, 0.1% SDS, 0.5% sodium deoxycholate, and 1x protease inhibitor) and sonicated (Bioruptor). Lysate was RNAse I (Ambion, 1:25) treated to fragment RNA. Protein-RNA complexes were immunoprecipitated using the rabbit polyclonal anti-Hnrnpa2b1 (ab31645, Abcam) antibody. One size-matched input (SMInput) library was generated per biological replicate using an identical procedure without immunoprecipitation. Stringent washes were performed as described, RNA was dephosphorylated (FastAP, Fermentas), T4 PNK (NEB), and a 3’ end RNA adaptor was ligated with T4 RNA ligase (NEB). Protein-RNA complexes were resolved on an SDS-PAGE gel, transferred to nitrocellulose membranes, and RNA was extracted from membrane. After RNA precipitation, RNA was reverse transcribed using SuperScript IV (Thermo Fisher Scientific), free primer was removed, and a 3’ DNA adapter was ligated onto cDNA products with T4 RNA ligase (NEB). Libraries were PCR amplified and dual-indexed (Illumina TruSeq HT). Pair-end sequencing was performed on Illumina NextSeq sequencer.

#### eCLIP bioinformatic and statistical analysis

Read processing and cluster analysis for Hnrnpa2b1 eCLIP was performed as previously described (Van Nostrand et al., 2016; Vogler et al., 2018). Read processing and cluster analysis for Hnrnpa2b1 eCLIP was performed as previously described. Briefly, 3’ barcodes and adapter sequences were removed using standard eCLIP scripts. Reads were trimmed, filtered for repetitive elements, and aligned to the mm9 reference sequence using STAR. PCR duplicate reads were removed based on the read start positions and randomer sequence. Bigwig files for genome browser display were generated based on the location of the second of the paired end reads. Peaks were identified using the encode_branch version of CLIPPER using the parameter “-s mm9.” Peaks were normalized against size matched input by calculating fold enrichment of reads in IP versus input and were designated as significant if the number of reads in the IP sample was greater than in the input sample, with a Bonferroni corrected Fisher exact p-value less than 10^−8^.

#### RNA-sequencing library preparation and sequencing

C2C12 Hnrnpa2b1-WT and KO myoblasts were seeded at 6 x 10^6^ cells per 15 cm plate, grown for 24hrs at 37C, 5% CO2 and differentiated via serum withdrawal and ITS supplementation for two days. Differentiated C2C12 myoblasts and myotubes were washed with PBS in a laminar flow hood and scraped from the tissue culture plate. Total RNA was extracted using a Qiagen RNeasy Plus Mini Kit, following manufacturer instructions. Isolated RNA quality was assessed by the CU Boulder BioFrontiers Next-Gen Sequencing core facility using an Agilent tape station. Isolated RNA was sent to the University of Colorado Cancer Center Genomics and Microarray Core for “Ribominus” Ribosomal RNA depletion, NGS library preparation and total RNA-sequencing.

#### RNA-sequencing informatics

All RNA sequencing data preprocessing was carried using standardized Nextflow RNA-sequencing (nf-core/rnaseq) with STAR alignment (genome GRCm39), transcriptome mapping with Salmon, and *in silico* ribosomal depletion using SortMeRNA. Differential gene expression was performed on feature counts tables using DESeq2 (Love et al., 2014). RNA splicing analysis was performed using Leafcutter (Li et al., 2018).

#### CLIP/splicing cluster analysis

The locations of splicing clusters generated by leafcutter and hnRNP A2/B1 eCLIP peaks from myotubes were each compared against genes in the mm10 refGene table from the UCSC Genome browser to determine the number of clusters or peaks overlapping each gene. The overlap of each cluster or peak was then subdivided into coverage of either UTR or segments of the gene, divided into percentiles along the gene. These were normalized and plotted against each other using R and ggplot2. (Script attached below, R command available on request)

#### Relative distance

Relative distance between splicing clusters generated by leafcutter and hnRNP A2/B1 eCLIP peaks from myotubes were generated using the bedtools reldist command, using eCLIP peaks as the first input, and splicing clusters in genes with eCLIP peaks as the second input (Favorov et al., 2012).

#### QAPA Poly-A Analysis

As previously described, RNA sequencing reads were trimmed, *in silico* ribodepleted, and reads were mapped to an mm10 3’UTR annotation file using salmon v0.13.1. Quantification of Alternative Polyadenylation (QAPA) was then used to estimate relative alternative 3’UTR isoform usage (Ha et al., 2018). Lengthening or shortening of a gene’s poly(A) tail was determined by first calculating a proximal poly(A) site usage (PPAU) defined as the percentage of reads mapping to the most proximal poly(A) site relative to reads mapping to the whole 3’UTR. △PPAU was then calculated as the median PPAU A2B1 KO – median PPAU WT (three replicates per condition). A gene with a △PPAU > 20 was defined as having a shortened poly(A) tail and a gene with a △PPAU < -20 was defined as having a lengthened poly(A) tail.

#### Hnrnpa2b1 immunoprecipitation and qRT-PCR

C2C12 Hnrnpa2b1-WT and KO myoblasts were seeded at 6 x 10^6^ cells per 15 cm plate, grown 24hrs at 37C, 5% CO2 and harvested as undifferentiated myoblasts. Myoblasts were lysed in a CHAPS-based buffer and pre-cleared using protein-A bound Dynabeads (Thermo, 10001D). Pre-cleared cell lysates were incubated with rabbit polyclonal anti-Hnrnpa2b1 antibody (ab31645, Abcam). Antibody-bound Hnrnpa2b1 was bound to protein-A Dynabeads overnight and magnetically isolated from the whole-cell lysate. Hnrnpa2b1-bound RNA was isolated from the Dynabead-Antibody-Hnrnpa2b1 complex via TRIZol RNA purification. cDNA libraries were created from purified RNA via oligo-DT priming and SuperScriptIII-enzyme reverse transcription. cDNA libraries were probed against Hnrnpa2b1-eCLIP hits chosen as a subset of the significantly enriched gene ontology terms via quantitative, real-time PCR. Primers used target GAPDH, Hnrnpa2b1, Prpf19, Snrnp70, Sfpq, Mbnl1, Hnrnpa3 and Mef2a. (Primer sequence in supplementary table 1). qRT-PCR was performed using SYBR-Green qRT-PCR reagent (BioRad) and fluorescent emission was measured using a BioRad CFX384 Real-Time PCR Detection system.

