## Supplementary Figures for "RNA-Binding Proteins Direct Myogenic Cell Fate Decisions"

1. Department of Biochemistry, University of Colorado Boulder, CO, USA
2. Medical Scientist Training Program, University of Colorado Anschutz Medical Campus Aurora, CO, USA
3. Department of Molecular, Cellular and Developmental Biology, University of Colorado Boulder, CO, USA
4. Molecular Biology Program and Department of Biochemistry and Molecular Genetics, University of Colorado Anschutz Medical Campus, Aurora, CO, USA.
5. University of Colorado School of Medicine, RNA Bioscience Initiative, University of Colorado Anschutz Medical Campus, Aurora, CO, USA.
6. Center for Personal and Dynamic Regulomes, Stanford University, CA, USA 7. Howard Hughes Medical Institute, Stanford University, CA, USA 8. Department of Neurology and Neurological Sciences, Stanford University School of Medicine, Stanford, CA, USA. 9. Paul F. Glenn Center for the Biology of Aging, Stanford University School of Medicine, Stanford, CA, USA. 10. Department of Molecular and Cell Biology, University of California, Berkeley, CA, USA
11. Center for Tissue Regeneration, Repair, and Restoration, Veterans Affairs Palo Alto Health Care System, Palo Alto, CA, USA.
12. Howard Hughes Medical Institute, University of Colorado, Boulder, CO, USA
13. Department of Pathology, Stanford University
14. Department of Neuropathology, Stanford University
15. These authors contributed equally
16. Co-senior author

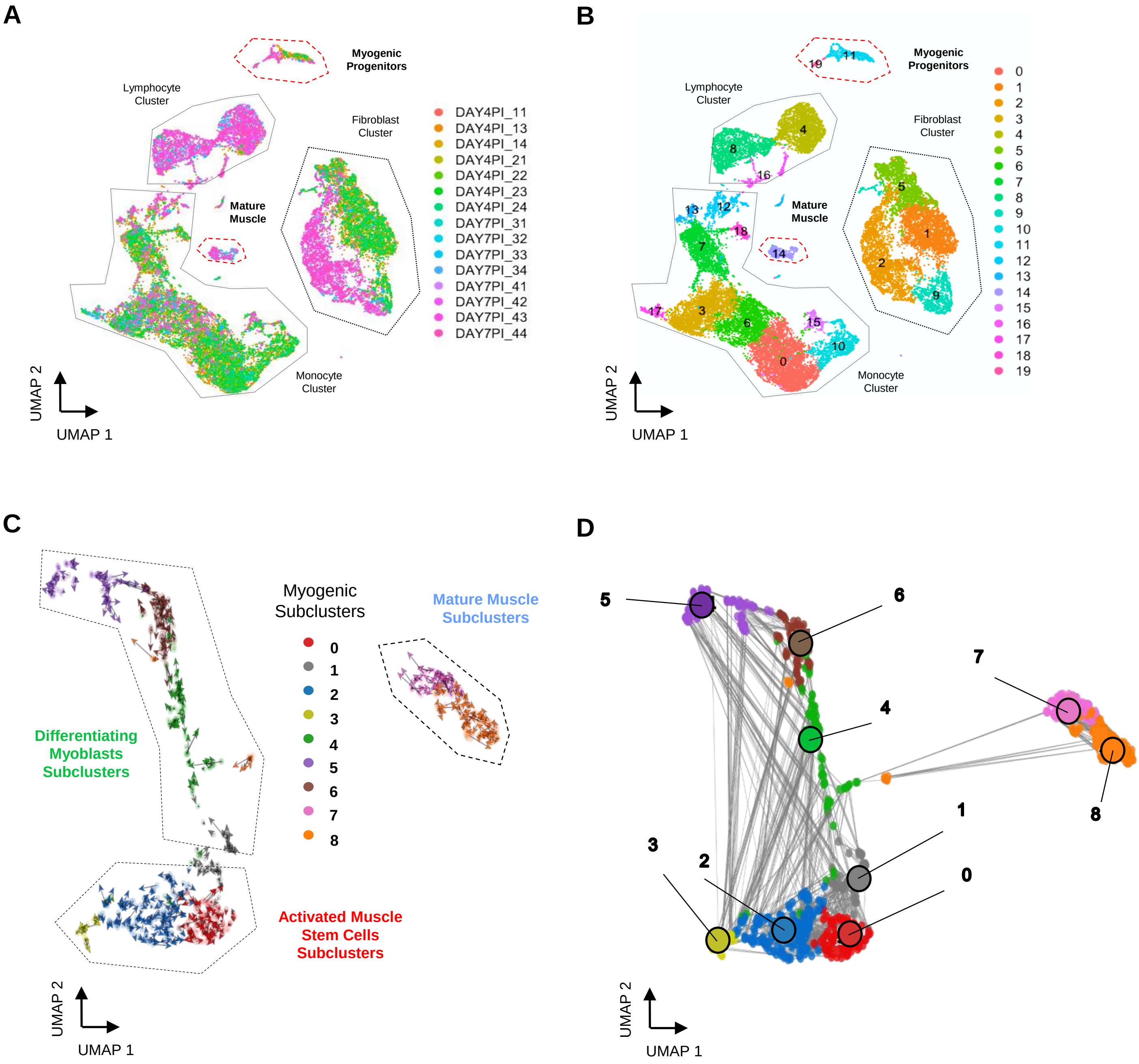


**Figure S1:** (Related to Figure 1) (A) UMAP of regenerating skeletal muscle at 4- and 7-dpi colored individual sequencing libraries (B) and by individual cellular clusters (C) RNA velocity of myogenic subclusters (D) Velocity inferred cell-cell connectivity between myogenic subclusters.

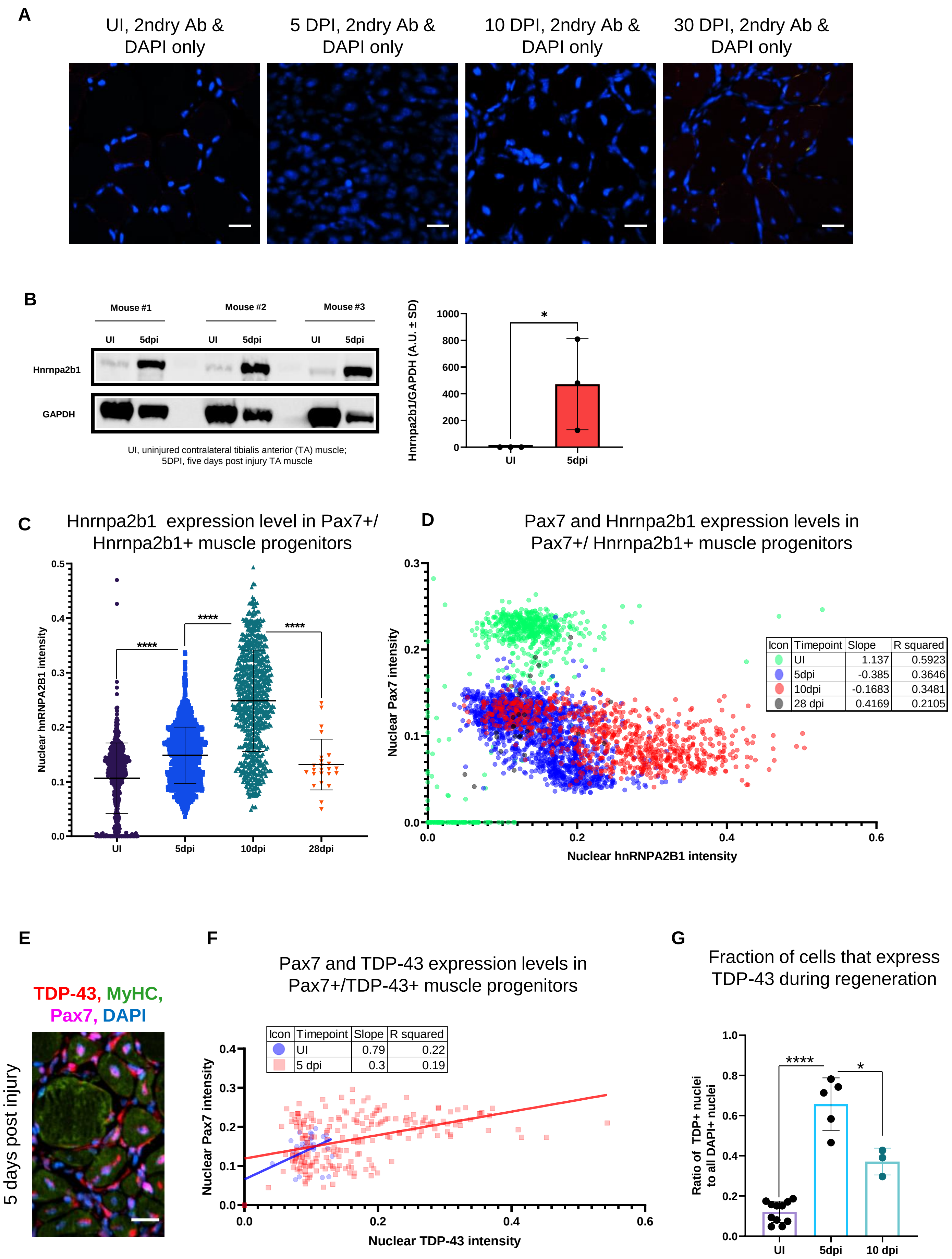
A
UI, 2ndry Ab & DAPI only
5 DPI, 2ndry Ab & DAPI only
10 DPI, 2ndry Ab & DAPI only
30 DPI, 2ndry Ab & DAPI only
B
Mouse #1
Mouse #2
Mouse #3
UI
5dpi
Hnrnpa2b1
GAPDH
UI, uninjured contralateral tibialis anterior (TA) muscle;
5DPI, five days post injury TA muscle
Hnrnpa2b1/GAPDH (A.U. ± SD)
1000
800
600
400
200
0
*
UI
5dpi
C
Hnrnpa2b1 expression level in Pax7+/ Hnrnpa2b1+ muscle progenitors
Nuclear hnRNPA2B1 intensity
0.5
0.4
0.3
0.2
0.1
0.0
****
****
****
UI
5dpi
10dpi
28dpi
D
Pax7 and Hnrnpa2b1 expression levels in Pax7+/ Hnrnpa2b1+ muscle progenitors
Nuclear Pax7 intensity
0.3
0.2
0.1
0.0
Nuclear hnRNPA2B1 intensity
0.0
0.2
0.4
0.6
Icon | Timepoint | Slope | R squared
UI | 1.137 | 0.5923
5dpi | -0.385 | 0.3646
10dpi | -0.1683 | 0.3481
28 dpi | 0.4169 | 0.2105
E
TDP-43, MyHC, Pax7, DAPI
5 days post injury
F
Pax7 and TDP-43 expression levels in Pax7+/TDP-43+ muscle progenitors
Icon | Timepoint | Slope | R squared
UI | 0.79 | 0.22
5 dpi | 0.3 | 0.19
Nuclear Pax7 intensity
0.4
0.3
0.2
0.1
0.0
Nuclear TDP-43 intensity
0.0
0.2
0.4
0.6
G
Fraction of cells that express TDP-43 during regeneration
Ratio of TDP+ nuclei to all DAPI+ nuclei
1.0
0.8
0.6
0.4
0.2
0.0
****
*
UI
5dpi
10 dpi

**Figure S3:** (Related to Figure 3) (A) Secondary only staining controls of muscle sections for all time points assessed by immunofluorescence (IF) in Figure 1A (B) Quantified western blot analysis of Hnrnpa2b1 protein expression from whole uninjured (UI) and 5-dpi (one-tail student t-test) (C) Hnrnpa2b1 nuclear intensity in Pax7-positive myonuclei in UI, 5-, 10-, 28-dpi regenerating muscle (D) Scatterplot and R-squared values for expression changes in myonuclear Pax7 and Hnrnap2b1 in UI, 5-, 10-, 28-dpi regenerating muscle (E) IF staining in 5-dpi regenerating muscle for Tardbp (TDP-43), Myosin heavy chain (MyHC), and Pax7 (F) Scatterplot and R-squared values for TDP-43 expression changes in Pax7 myonuclei in UI and 5dpi (G) TDP-43 nuclear expression UI, 5dpi, 10dpi. All images represent n=3 biological replicates; scale = 20µM. All quantified data represent mean ± SD, two-tail student t-test p-value: *=<0.05, ** =<0.01, ***=<0.001, ****=<0.0001 unless otherwise stated.

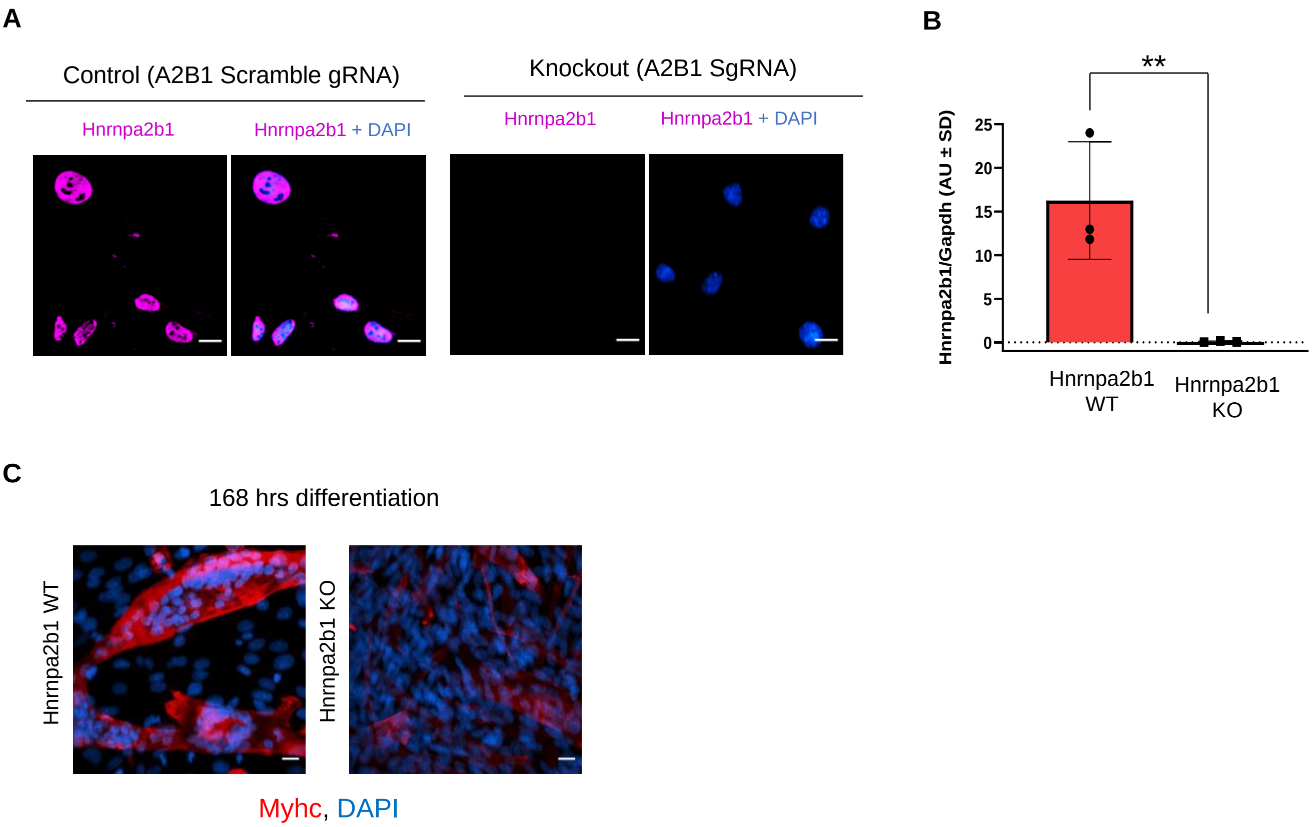


**Figure S4:** (Related to Figure 4) (A) Hnrnpa2b1 expression in WT and Hnrnpa2b1 KO myoblasts (scale = 10uM) (B) Western blot (WB) quantification of Hnrnpa2b1 in WT and Hnrnpa2b1 KO myoblasts (C) MyHC expression in WT and Hnrnpa2b1 KO 168 hour differentiated myotubes. All images represent n=3 biological replicates from 3 independent WT and KO Hnrnpa2b1 clones; scale = 20uM. All quantified data represent mean ± SD, two-tail student t-test p-value: **=<0.01.

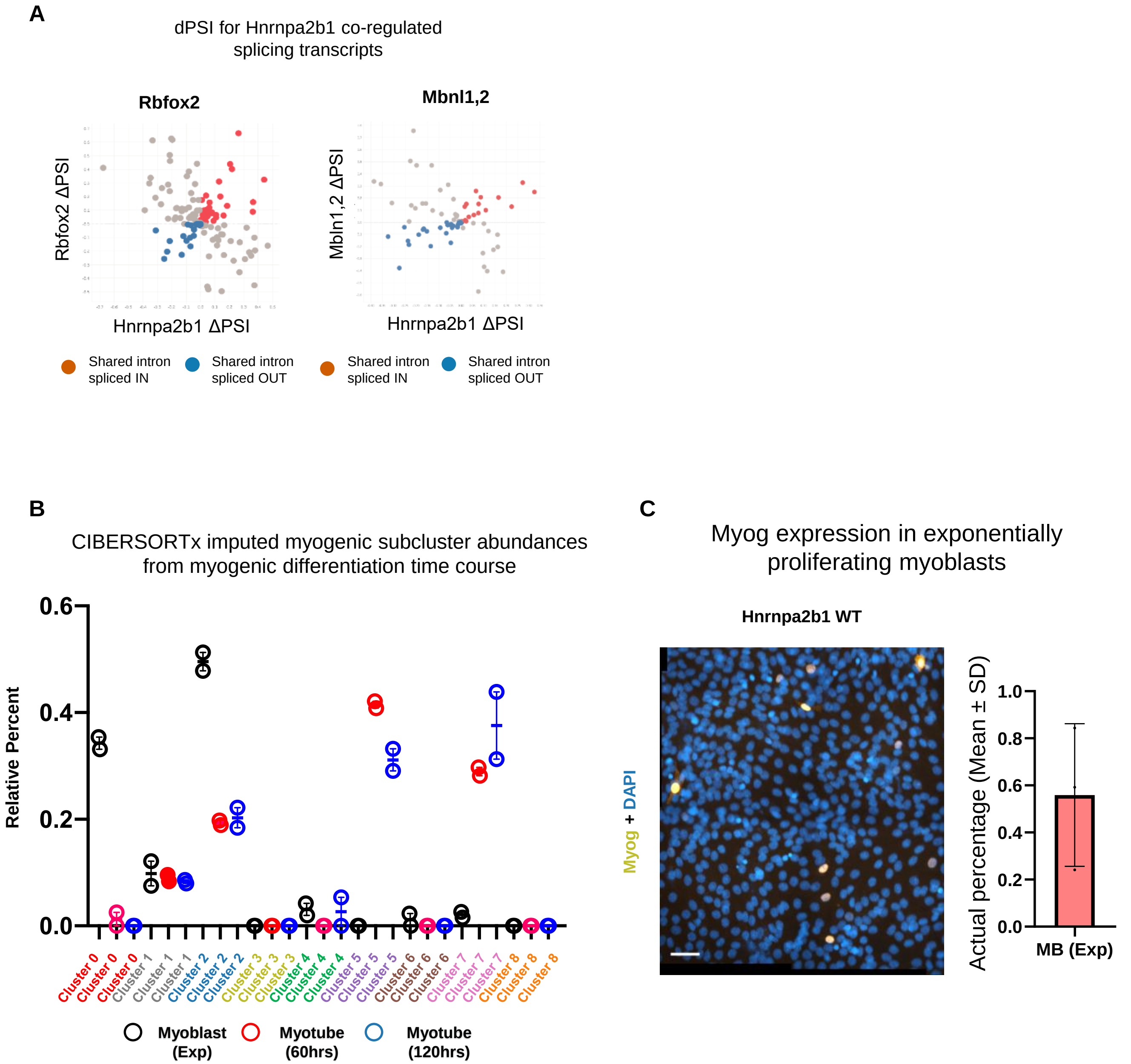


**Figure S5:** (Related to Figure 5) (A) Hnrnpa2b1-Rbfox or Hnrnpa2b1-Mbnl1/Mbnl2 dPSI (percent spliced in) changes in shared significantly altered spliced RNAs. (B) Myogenic-trained CIBERSORTx imputed percentages of myogenic clusters from RNA sequencing performed in biological duplicate of C2C12 differentiation time course (Trapnell et al., 2010). (C) Myogenin immunoreactivity and nuclear staining of exponentially proliferating *in vitro* myoblasts. Bar plot represents actual percentage of myogenin-expressing proliferating myoblasts, n=3 biological replicates, scale = 50uM.

### Figure S6

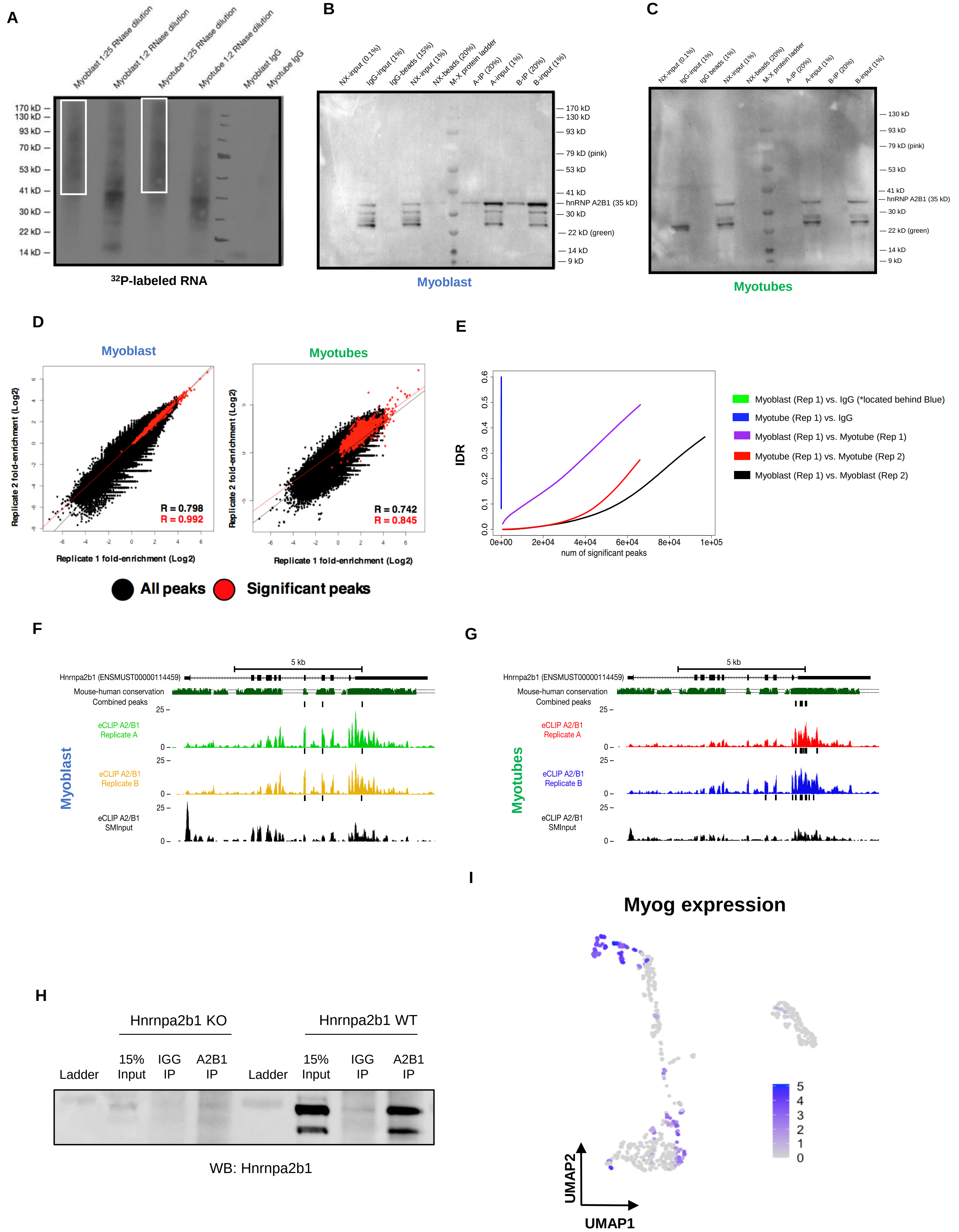

A
Myoblast 1:25 RNase dilution
Myoblast 1:2 RNase dilution
Myotube 1:25 RNase dilution
Myotube 1:2 RNase dilution
Myoblast IgG
Myotube IgG
170 kD
130 kD
93 kD
70 kD
53 kD
41 kD
30 kD
22 kD
14 kD
32P-labeled RNA
B
NX-input (0.1%)
IgG-input (1%)
IgG-beads (15%)
NX-input (1%)
NX-beads (20%)
M-X protein ladder
A-IP (20%)
A-input (1%)
B-IP (20%)
B-input (1%)
170 kD
130 kD
93 kD
79 kD (pink)
53 kD
41 kD
hnRNP A2B1 (35 kD)
30 kD
22 kD (green)
14 kD
9 kD
Myoblast
C
NX-input (0.1%)
IgG-input (1%)
IgG beads (1%)
NX-input (1%)
NX-beads (20%)
M-X protein ladder
A-IP (20%)
A-input (1%)
B-IP (20%)
B-input (1%)
130 kD
93 kD
79 kD (pink)
53 kD
41 kD
hnRNP A2B1 (35 kD)
30 kD
22 kD (green)
14 kD
9 kD
Myotubes
D
Myoblast
Myotubes
Replicate 2 fold-enrichment (Log2)
Replicate 1 fold-enrichment (Log2)
R = 0.798
R = 0.992
R = 0.742
R = 0.845
All peaks
Significant peaks
E
IDR
num of significant peaks
Myoblast (Rep 1) vs. IgG (*located behind Blue)
Myotube (Rep 1) vs. IgG
Myoblast (Rep 1) vs. Myotube (Rep 1)
Myotube (Rep 1) vs. Myotube (Rep 2)
Myoblast (Rep 1) vs. Myoblast (Rep 2)
F
5 kb
Hnrnpa2b1 (ENSMUST00000114459)
Mouse-human conservation
Combined peaks
eCLIP A2/B1 Replicate A
eCLIP A2/B1 Replicate B
eCLIP A2/B1 SMInput
Myoblast
G
5 kb
Hnrnpa2b1 (ENSMUST00000114459)
Mouse-human conservation
Combined peaks
eCLIP A2/B1 Replicate A
eCLIP A2/B1 Replicate B
eCLIP A2/B1 SMInput
Myotubes
H
Hnrnpa2b1 KO
Hnrnpa2b1 WT
Ladder
15% Input
IGG IP
A2B1 IP
WB: Hnrnpa2b1
I
Myog expression
UMAP1
UMAP2

**Figure S6:** (Related to Figure 6) (A) Autoradiogram of 32P-labelled Hnrnpa2b1 RNA –RNA complexes fractionated by PAGE (B) and (C) Immunoprecipitation of Hnrnpa2b1 RNA complexes used for eCLIP in C2C12 myoblasts or myotubes (n = 2 biologically independent samples) (D) Scatter plots indicate correlation between significant Hnrnpa2b1 RNA eCLIP peaks in biological replicates. Scatter plots represent fold enrichment for each region in Hnrnpa2b1 RNA eCLIP relative to paired size-matched input with significant peaks in red ($P \leq 10^{-8}$ over size-matched input) (E) Irreproducible discovery rate (IDR) analysis comparing peak fold enrichment across indicated eCLIP datasets (F) IGV tracing showing Hnrnpa2b1 RNA 3'UTR eCLIP peaks in Hnrnpa2b1 RNA transcript in myoblasts and (G) myotubes (H) Western blot for Hnrnpa2b1 used in RIP experiments in both Hnrnpa2b1 RNA wildtype and KO C2C12 cells (I) UMAP displaying Myog expression across myogenic single cells.
